# Functional metagenomics using *Pseudomonas putida* expands the known diversity of polyhydroxyalkanoate synthases and enables the production of novel polyhydroxyalkanoate copolymers

**DOI:** 10.1101/042705

**Authors:** Jiujun Cheng, Trevor C. Charles

## Abstract

Bacterially produced biodegradable polyhydroxyalkanoates with versatile properties can be achieved using different PHA synthase enzymes. This work aims to expand the diversity of known PHA synthases via functional metagenomics, and demonstrates the use of these novel enzymes in PHA production. Complementation of a PHA synthesis deficient *Pseudomonas putida* strain with a soil metagenomic cosmid library retrieved 27 clones expressing either Class I, Class II or unclassified PHA synthases, and many did not have close sequence matches to known PHA synthases. The composition of PHA produced by these clones was dependent on both the supplied growth substrates and the nature of the PHA synthase, with various combinations of SCL-and MCL-PHA. These data demonstrate the ability to isolate diverse genes for PHA synthesis by functional metagenomics, and their use for the production of a variety of PHA polymer and copolymer mixtures.

## Introduction

Polyhydroxyalkanoates (PHAs) are natural polyesters biosynthesized by a variety of bacteria under unbalanced growth conditions. They serve as reserves of carbon and reducing power and aid in survival during starvation or stress conditions (Verlinden et al. 2007). These biodegradable and environmentally friendly polymers can be used as alternative materials to conventional petrochemical-based plastics. PHAs are classified into short-chain-length (SCL, C3-C5) and medium-chain-length (MCL, ≥C6), and copolymers (SCL + MCL) based on the number of carbon atoms per monomer. MCL-PHA is generally more useful than SCL-PHA due to it being less brittle and more flexible. To produce PHA with versatile properties cost-effectively, strategies have involved mining new PHA synthase enzymes (PhaC), engineering PhaC proteins and modifying the metabolic pathways of the production strains (Keshavarz and Roy 2010; Schallmey et al. 2011; Cheema et al. 2012; Park et al. 2012; Tripathi et al. 2013; Meng et al. 2014).

PHA molecules consist of over 150 possible constituent monomers of hydroxyl fatty acids (Meng et al. 2014) (Steinbüchel and Lütke-Eversloh 2003). PHA synthase enzymes catalyze the joining of the hydroxyl group of one monomer with the carboxyl group of another by an ester bond to form PHA polymers. The monomeric composition of PHA is determined primarily by PhaC, although the available carbon source and metabolic pathways also influence the properties of PHA (Verlinden et al. 2007). PhaC proteins are grouped into four classes based on amino acid sequence, substrate specificity and subunit composition (Rehm 2003). Class I and II PhaC consist of one subunit (PhaC), but those in Class III and IV are composed of two subunits (PhaC+PhaE and PhaC+PhaR, respectively). Class I and IV PhaCs synthesize SCL-PHA whereas Class II polymerize MCL-PHA. Class III PhaCs can synthesize both SCL and MCL monomers.

The key Class I PHA synthesis pathway genes include *phaC* (or *phbC*), *phaA* (or *phbA*) and *phaB* (or *phbB*). The three genes are often, but not always, clustered in a single operon. The acetoacetyl-CoA reductase (EC 2.3.1.9) encoded by the *phaA* and β-ketothiolase (EC 1.1.1.36) encoded by the *phaB* convert acetyl-CoA to (R)-3-hydroxybutyryl-CoA (3HB-CoA) through acetoacetyl-CoA. The Class-I PhaC (EC 2.3.1.-) then polymerizes 3HB-CoA into polyhydroxybutyrate (PHB) (Rehm 2003). The Class II *pha* cluster is well conserved in *Pseudomonas* and consists of two PHA synthase genes (*phaC1* and *phaC2*) flanking a PHA depolymerase gene (*phaZ*), and *phaD* encoding a transcriptional activator of *pha* genes. Phasin-encoding *phaF* and *phaI* gene are transcribed divergently to other *pha* genes. MCL monomers ((R)-3-hydroxylacyl-CoA) are derived from β-oxidation of fatty acids or fatty acid *de novo* synthesis from unrelated carbon sources (Tortajada et al. 2013).

Traditional culture-based strategies for obtaining new biocatalysts are limited by the inability to cultivate the majority of environmental microbes. While sequence-based metagenomics can identify genes homologous to those present in available sequence databases, it is difficult to reliably predict the function of truly new genes through homology-based analysis. In contrast, functional metagenomics involves the construction of gene libraries from microbial community genomic DNA and functional screening for novel enzymes of interest or potential industrial applications (Simon and Daniel 2011). Functional metagenomics has the potential to identify truly novel sequences for a given function.

In previous work, functional metagenomics was used to isolate new Class I PhaC from soil metagenomic clones by Nile red staining and phenotypic screening in *α-Proteobacteria Sinorhizobium meliloti* (Schallmey et al. 2011). In another study, *phaC* genes encoding both Class I and II PhaC proteins were PCR amplified from oil-contaminated soil library clones (Cheema et al. 2012). One of the isolated genes produced PHA copolymer when expressed in *Pseudomonas putida*. Moreover, partial *phaC* genes were also obtained from metagenomic DNA via direct PCR amplification (Foong et al. 2014; Pärnänen et al. 2015; Tai et al. 2015). Building on these previous studies, we constructed a PHA^-^ strain of *P. putida* KT2440 to use as a surrogate host and functionally identified *phaC* genes after screening millions of agricultural wheat soil metagenomic clones. A total of 27 PHA^+^ clones were obtained. Accumulation and monomer composition of PHA directed by seven of these clones were further examined.

## Methods

### Bacterial strains, plasmids, cosmids and growth conditions

Bacterial strains, plasmids and cosmids are listed in Table 1. *E. coli* strains were grown at 37°C in LB (Lennox) medium (1% tryptone (w/v), 0.5% yeast extract (w/v), and 0.5% NaCl (w/v), pH 7). *Pseudomonas* strains were grown at 30°C in LB or 0.1N M63 minimal medium (Escapa et al. 2011) supplemented with 0.5% sodium octanoate (w/v), 0.5% nonanoic acid (v/v) or 1% gluconic acid (w/v). *S. meliloti* was grown in LB or YM medium (Schallmey et al. 2011). Antibiotics were used at the following concentrations: streptomycin, 200 μg/ml for *S. meliloti* and 100 μg/ml for *E. coli*; kanamycin, 100 μg/ml for *Pseudomonas* and 50 μg/ml for *E. coli*; neomycin, 200 μg/ml; rifampicin, 100 μg/ml; gentamicin, 10 μg/ml for *E. coli* and 100 for *P. putida*; and tetracycline 20 μg/ml for *E. coli* or 40 μg/ml for *P. putida*.

**Table 1.**
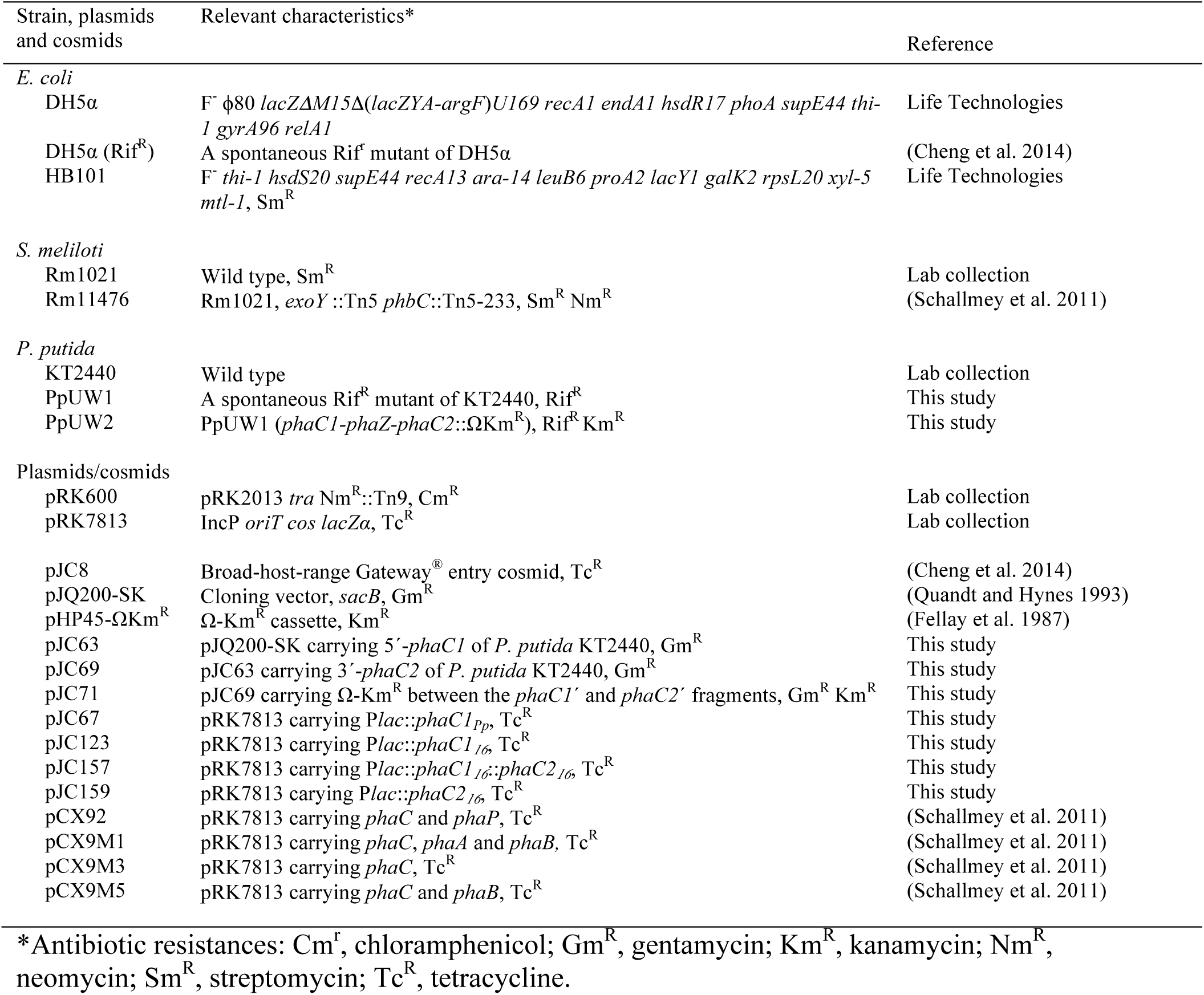
Bacterial strains, plasmids and cosmids

### Construction of PHA^-^ strain PpUW2

DNA oligonucleotides are listed in Table 2. A Rif^R^ spontaneous mutant PpUW1 of *P. putida* KT2440 was generated by plating a culture of strain KT2440 on a LB Rif plate, followed by single colony purification. To construct the PHA deficient strain PpUW2 (*phaC1ZC2*) of *P. putida* PpUW1, a 943-bp DNA fragment containing 5′-*phaC1* region (766 bp) was PCR amplified using *P. putida* KT2440 genomic DNA as a template and primer pair JC161-JC162, digested with HindIII-BamHI and then cloned into the same sites in pJQ200-SK (Quandt and Hynes 1993), yielding plasmid pJC63. Another 876-bp DNA fragment containing the 3′-*phaC2* gene (120 bp) was PCR amplified using primers JC163-JC164, digested with BamHI-SalI and then cloned into the same sites in pJC63 to obtain plasmid pJC69. An omega-Km cassette was obtained from pHP45Ω-Km (Fellay et al. 1987) by BamHI digestion and then inserted into the same site in pJC69 to obtain pJC71. Plasmid pJC71 was then conjugated into *P. putida* PpUW1 in a triparental mating using helper plasmid pRK600. Single cross-over recombination of pJC71 into the *P. putida* chromosome was selected with Rif and Gm. A double cross-over event was achieved by growing a single Rif^R^ Gm^R^ colony overnight in LB, making serial dilutions and then spreading on LB Km with 5% sucrose. The resulting PHA^-^ strain PpUW2 was verified by examining Gm sensitivity (lost plasmid backbone) and absence of PHA production in LB supplemented with 0.5% octanoate (w/v), and was further confirmed by PCR amplification analysis.

**Table 2.**
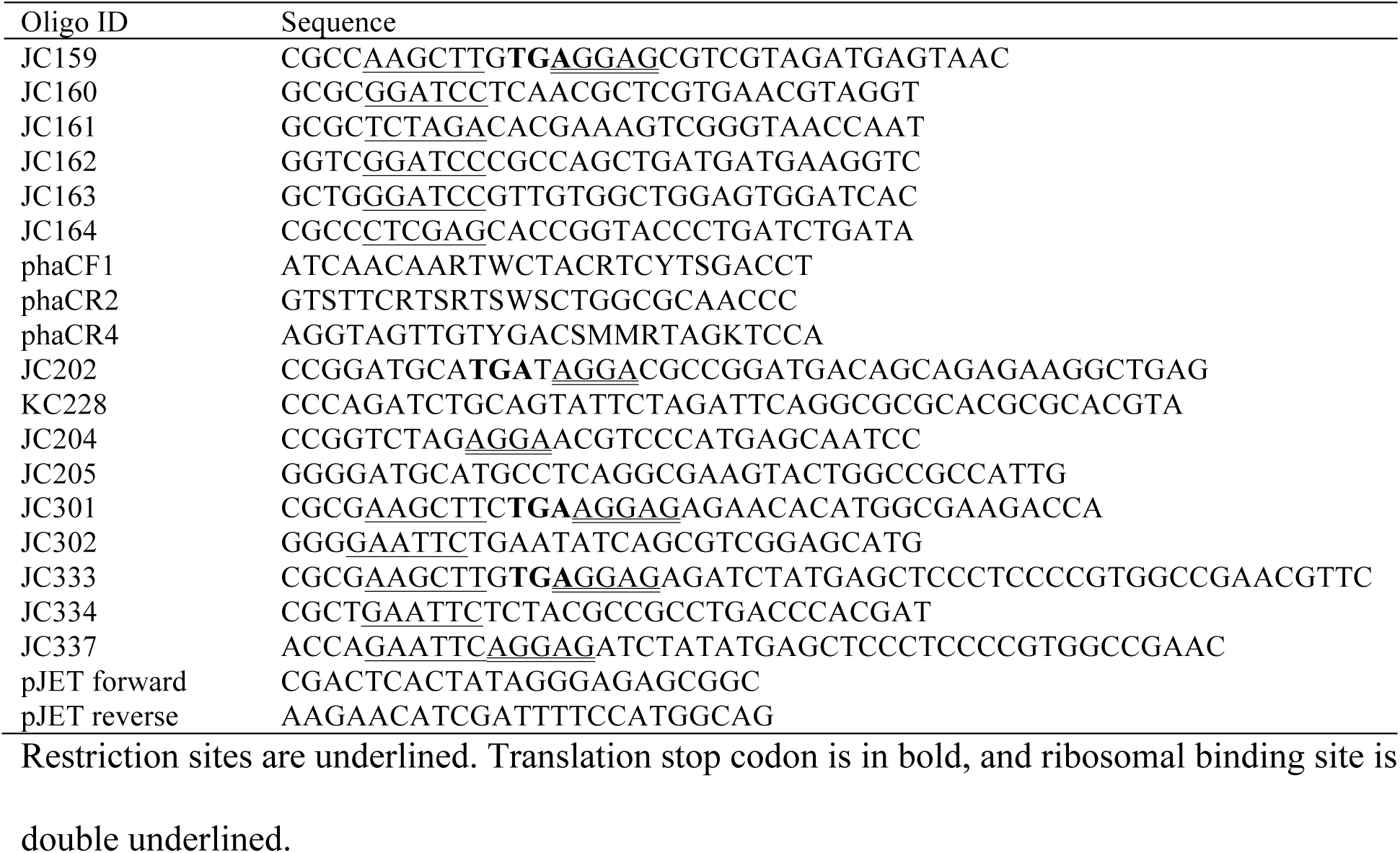
DNA oligonucleotides

### Phenotypic screening for *phaC* genes from metagenomic library clones

Construction of the metagenomic DNA library (11AW) of agricultural wheat soil was described previously (Cheng et al. 2014). The 11AW library contains 9 × 10^6^ clones hosted in *E. coli* HB101. The pooled library clones (0.5 ml of 300 ml stock) were conjugated *en masse* into *P. putida* PpUW2 (PHA^-^) with the helper plasmid pRK600. Mating mixture was diluted serially with 0.85% NaCl and ~20,000 transconjugants recovered on each LB Tc plate (15 mm × 150 mm) supplemented with 0.5% Na octanoate (w/v). The plates were incubated at 30°C for 24 h and then kept at 22°C for 2-6 days. Potential PHA^+^ clones of opaque white colour were streak purified on the LB octanoate plates, and verified on 0.1N M63 minimal medium plates (Escapa et al. 2011) supplemented with 0.5% octanoate (w/v) and Nile red (0.5 μg/ml). PHA^+^ cosmids were transferred from *P. putida* to *E. coli* DH5α(Rif^R^), mobilized by pRK600. The cosmid DNA was isolated from *E. coli* strains using a GeneJET Plasmid Miniprep Kit (Thermo Scientific), digested by EcoRI-HindIII-BamHI (Thermo Scientific), and then resolved on 1% TAE agarose gels. Cosmids with distinct electrophoretic patterns were conjugated back to *P. putida* PpUW2 (PHA^-^) to confirm their PHA^+^ phenotype.

### Complementation of *S. meliloti* (*phbC*)

11AW cosmid DNA encoding Class I and II *phaC* genes was introduced into *S. meliloti* Rm11476 (Schallmey et al. 2011) via triparental conjugation. Transconjugants were selected on LB SmNmTc plates, and then streaked on YM plates containing Nile red (Schallmey et al. 2011)for visualizing PHB production.

### DNA sequencing and bioinformatics

KOD Xtreme DNA polymerase (Novagen) was used for all PCR. Primers are listed in Table 2. PCR reactions consisted of one cycle of 94°C for 5 min, 30 cycles of 94°C for 30 s, 53°C for 30 s and 68°C for 30 s, and final extension at 68°C for 10 min. The internal regions of *phaC* genes were PCR amplified from PHA^+^ cosmids using primer phaCF1 and phaCR4 (Sheu et al. 2000). When no PCR product of correct size was obtained, a semi-nested PCR was performed with the primers phaCF2 and phaCR4 as described previously (Sheu et al. 2000). PCR products were resolved on 2% agarose gels, isolated from the gels using EZ-10 Spin Column DNA Gel Extraction Kit (Bio Basic), and then cloned into pJET1.2 vector (Thermo Scientific). Sequences of the cloned partial *phaC* genes were obtained using pJET1.2 sequencing primers (Table 2).

End sequences of PHA^+^ cosmid DNA were obtained by Sanger sequencing using universal M13F and M13R primers. For high throughput sequencing, 68 cosmids of 11AW PHA^+^ clones and 28 cosmids from other research projects were grouped into 24 pools. Tagmentation of pooled cosmid DNA, PCR amplification and clean-up, and library normalization were performed with the Nextera XT DNA library and index kits (Illumina), according to the supplier’s recommendation. The library was sequenced using MiSeq 500-cycle version 2 reagent (Illumina). DNA sequences were assembled using SPAdes Genome Assembler 3.5.0 (BaseSpace, Illumina). Cosmid clones were identified based on the available Sanger end sequences (Lam et al. 2014) and ORFs were annotated with MetaGeneMark (Zhu et al. 2010). Predicted protein sequences were analyzed by BLAST against the non-redundant protein databases (NCBI). Multiple-sequence alignment was performed using MUSCLE (Edgar 2004). Phylogenetic analysis was performed using MEGA6 (Tamura et al. 2013). Origin of cloned metagenomic DNA was predicted by PhyloPythia (Patil et al. 2012).

### Cloning of *phaC1*_*Pp*_ gene of *P. putida* KT2440

To construct a *phaC* expression cosmid for a positive control, the *phaC1*_*Pp*_ gene (*Pp_5003*) of *P. putida* KT2440 was PCR amplified using primers JC159-JC160 and cloned into the HindIII and BamHI sites in the broad-host-range vector pRK7813 (Cheng et al. 2014), to obtain construct pJC67 (P*lac*∷*phaC1*_*Pp*_). A stop codon (TGA) for terminating the translation of *lacZα* upstream of the *phaC1*_*Pp*_, and a ribosome-binding site (AGGAG) were incorporated into the primer JC159 (Table 2).

### Subcloning *phaC* genes of 11AW metagenomic clone 16

The Class II *phaC1*_*16*_ gene of 11AW clone 16 was PCR amplified with oligos JC301 and JC302, and cloned into the HindIII-EcoRI sites in pRK7813 to obtain pJC123. The *phaC2*_*16*_ was amplified using primers JC337 and JC334, and inserted into the EcoRI site in pJC123, yielding pJC157. The orientation of cloned *phaC2*_*16*_ was verified by restriction enzyme mapping. The *phaC2*_*16*_ gene was also obtained by PCR using oligos JC333 and JC334, and cloned into the HindIII-EcoRI sites in pRK7813 to yield pJC159. A stop codon (TGA) for terminating the translation of *lacZα* gene and ribosome-binding sites were added in primers JC333 and JC337. The cloned *phaC* genes were verified by DNA sequencing.

### Estimation of PHA production by fluorescent spectrometry

A modification of the previously described method (Wu et al. 2003) was used. *S. meliloti* cells (2.5%, v/v) were added to YM medium. All cultures were incubated at 30°C and 200 rpm for 48-72 h. The OD_600_ values of 200 μl cultures were measured in 96-well microtiter plates with the Multiskan Spectrum spectrophotometer (Thermo Labsystems). Samples (180 μl) were stained with 20 μl of Nile red (2.5 μg/ml) in the dark for 1 h. Fluorescent intensity was measured at excitation (485 nm) and emission (595 nm) with the FilterMax F5 Multi-mode microplate reader (Molecular Devices). PHA content was calculated based on the equation (fluorescent intensity/OD_600_×0.9).

### PHA production, extraction and characterization

*P. putida* PpUW1 (PHA^+^) or PpUW2 (PHA^-^) carrying PHA^+^ clones were grown overnight in LB with or without Tc, washed once with 0.85% NaCl, and then subcultured (2%, v/v) in the 0.1N M63 medium with or without Tc, supplemented with 0.5% Na octanoate, 0.5% nonanoic acid or 1% gluconic acid. The cultures were grown at 30°C and at 200 rpm for 48 hrs. Cells were collected by centrifugation at 20°C and at 9000 × *g* for 15 min, washed once with deionized water. Cell dry weight (CDW) was obtained after drying the cells at 95°C for 24 hr. For PHA methanolysis, cell pellet (~15 mg) was suspended in 2 ml of chloroform and 2 ml of methanol with 15% H_2_SO_4_ (v/v), and incubated at 100°C for 5 h. After the reaction mixture was cooled down to 20°C, 1 ml of deionized water was added, and vortexed vigorously for 1 min. The chloroform phase was passed through a cotton plug in a Pasteur glass pipette to remove any cell debris. Methanolyzed sample (1 μl) was analyzed with GC-MS (an Agilent 7975B GC equipped with Agilent 5975B inert XL EI/CI MSD and an HP-5MS capillary column). Oven temperature was run at initial 50°C for 5 min with a ramp of 20°C/min to 280°C, and then held for 10 min. The flow rate of Helium carrier gas was 1.2 ml/min. Methylated PHA monomers were identified using the Agilent enhanced MSD chemstation (E.02.01.117). PHA standards were kindly provided by Dr. Bruce A. Ramsay (Polyferm Canada).

## Results

### Isolation of metagenomic clones for PHA production

To employ *P. putida* as a surrogate host to isolate 11AW (agricultural wheat soil) metagenomic library clones encoding functional PhaC, it was first necessary to construct a PHA synthesis mutant. Expression of the contiguous three genes encoding PHA synthase PhaC1_Pp_(Pp_5003), PHA depolymerase PhaZ_Pp_ (Pp_5004) and PHA synthase PhaC2_Pp_ (Pp_5005) in *P. putida* KT2440 Rif^R^ derivative PpUW1 was disrupted by deletion of 3437-bp comprised of the 3′ region (911 bp) of *phaC1*_*Pp*_ (1677 bp), *phaZ*_*Pp*_ and the 5′ region (1560 bp) of *phaC2*_*Pp*_ (1680 bp) and replacement with an omega-Km kanamycin resistance insert, resulting in the PHA^-^ strain PpUW2. Transfer of the negative control cosmid vector pJC8 to *P. putida* PpUW2 did not result in detectable PHA production. However, introduction of pJC67 (P*lac*∷*phaC1*_*Pp*_) *in trans* restored the PHA^+^ phenotype (Fig. 1). These data suggested that the strain PpUW2 could be used for screening of PhaC-encoding metagenomic clones. The 11AW library clones (Cheng et al. 2014) were transferred to *P. putida* PpUW2 via en masse triparental conjugation. Following selection of *P. putida* PpUW2 transconjugants (~4 million) on LB Km Tc plates supplemented with 0.5% Na octanoate, we obtained 72 clones that exhibited greater opacity than the PpUW2 PHA^-^ recipient strain. The PHA-producing phenotype of those clones was verified by visualizing the fluorescence of Nile red-stained PHA in 0.1N M63 minimal medium (Escapa et al. 2011) supplemented with Na octanoate as the sole carbon source (Fig. 1). Restriction digest of DNA from the 72 clones demonstrated 68 distinct restriction patterns, suggesting the presence of a broad diversity of DNA origin in those PHA^+^ clones.

**Fig. 1.**
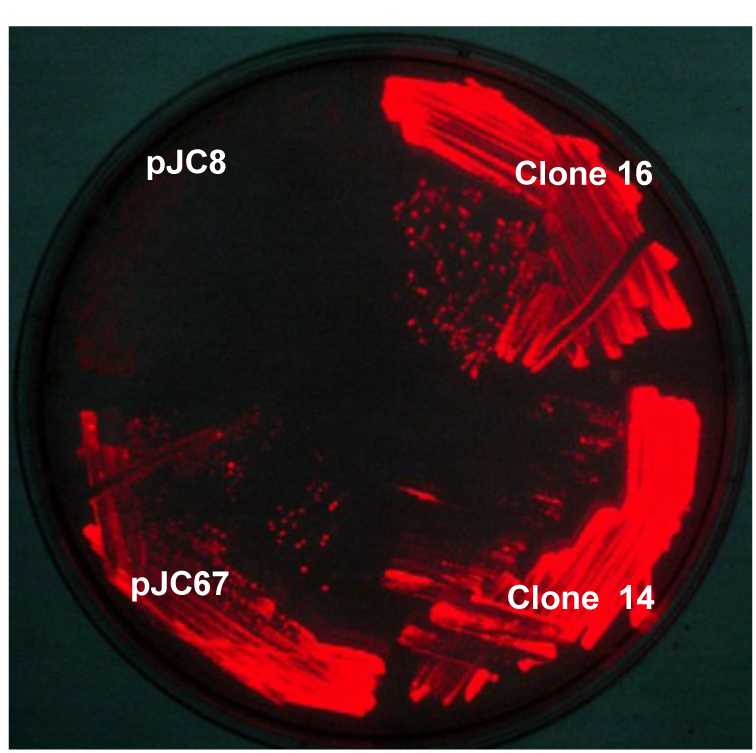
Activity-based screening of 11AW metagenomic library clones carrying PHA biosynthesis genes in *P. putida* PpUW2 (PHA^-^). PpUW2 recombinant strains were grown in 0.1N M63 medium containing 0.5% Na octanoate and Nile red (0.5 μg/ml). Cosmid pJC8 and pJC67 (P*lac*∷*phaC1*_*Pp*_) were used as negative and positive controls respectively. 11AW clones 14 and 16 were streaked on the same selection plate to verify the PHA^+^ phenotype initially screened on LB with 0.5% (w/v) octanoate.

### Identification of *phaC* genes encoding PHA synthases

Internal regions (~500 bp) of the *phaC* genes of 18 distinct cosmid clones were initially obtained by PCR amplification using degenerate primers PhaCF1 and PhaCR4 (Sheu et al. 2000). For additional 4 clones, *phaC* fragments (PhaC_3_, PhaC_7_, PhaC_10_, and PhaC_15_) of ~400 bp were generated by nested-PCR using primer pair PhaCF2-PhaCR4 as described previously (Sheu et al. 2000). BLASTP analysis of the cloned gene products indicated that 13 PhaC proteins could be grouped into Class I while the other 9 were categorized as Class II PHA synthases.

To identify all *pha* genes on the isolated clones, DNA sequences were obtained by high throughput sequencing, and additional *pha* gene loci were identified in partially and fully assembled clones (Table 3; Supplementary 1). The identified PHA synthases could be classified into 3 groups based on amino acid sequences (Fig. 2). Class I *phaC* genes were annotated in 17 clones. The *phaC* and *phaAB* genes were adjacently located in 12 of these clones, whereas the *phaB* gene was located distantly downstream of *phaCA* genes in clone 25 and clone P1N3 (Supplementary 1). A *phaR* gene was located immediately downstream of *phaCB* genes in partially assembled clone P11N2. The *phaC* and *phaB* genes flanked a *phaZ* gene in the partially assembled clone P2N8. The metagenomic DNA in these Class I clones was predicted to originate from *Gemmatimonas*, *α-Proteobacteria* (*Sphingomonadaceae*), *β-Proteobacteria* (*Leptothrix, Rubrivivax, Janthinobacterium*) and γ-*Proteobacteria* (*Xanthomonadaceae*) (Table 3).

**Table 3.**
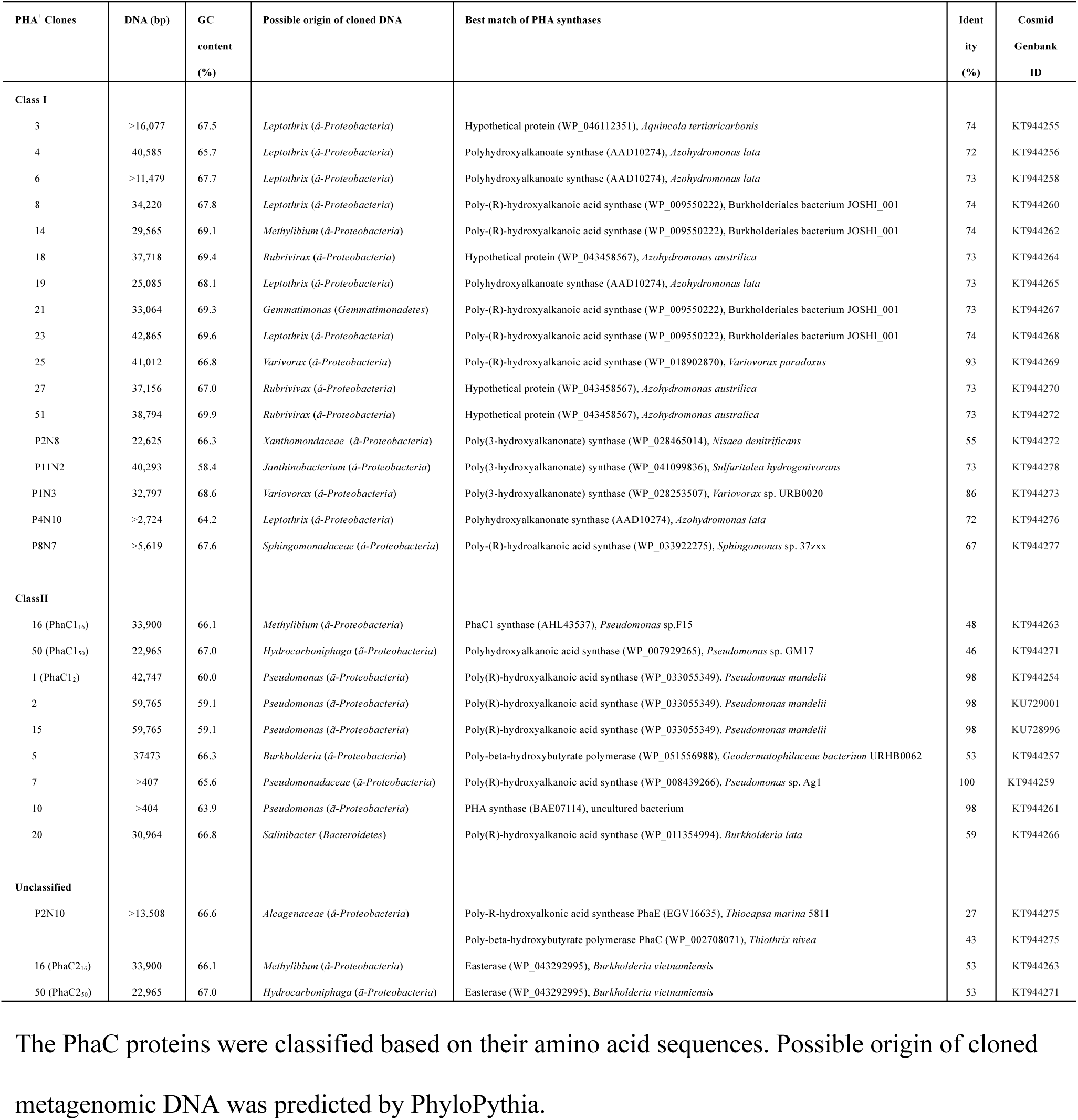
11AW metagenomic clones encoding PHA synthases (PhaC).

**Fig. 2.**
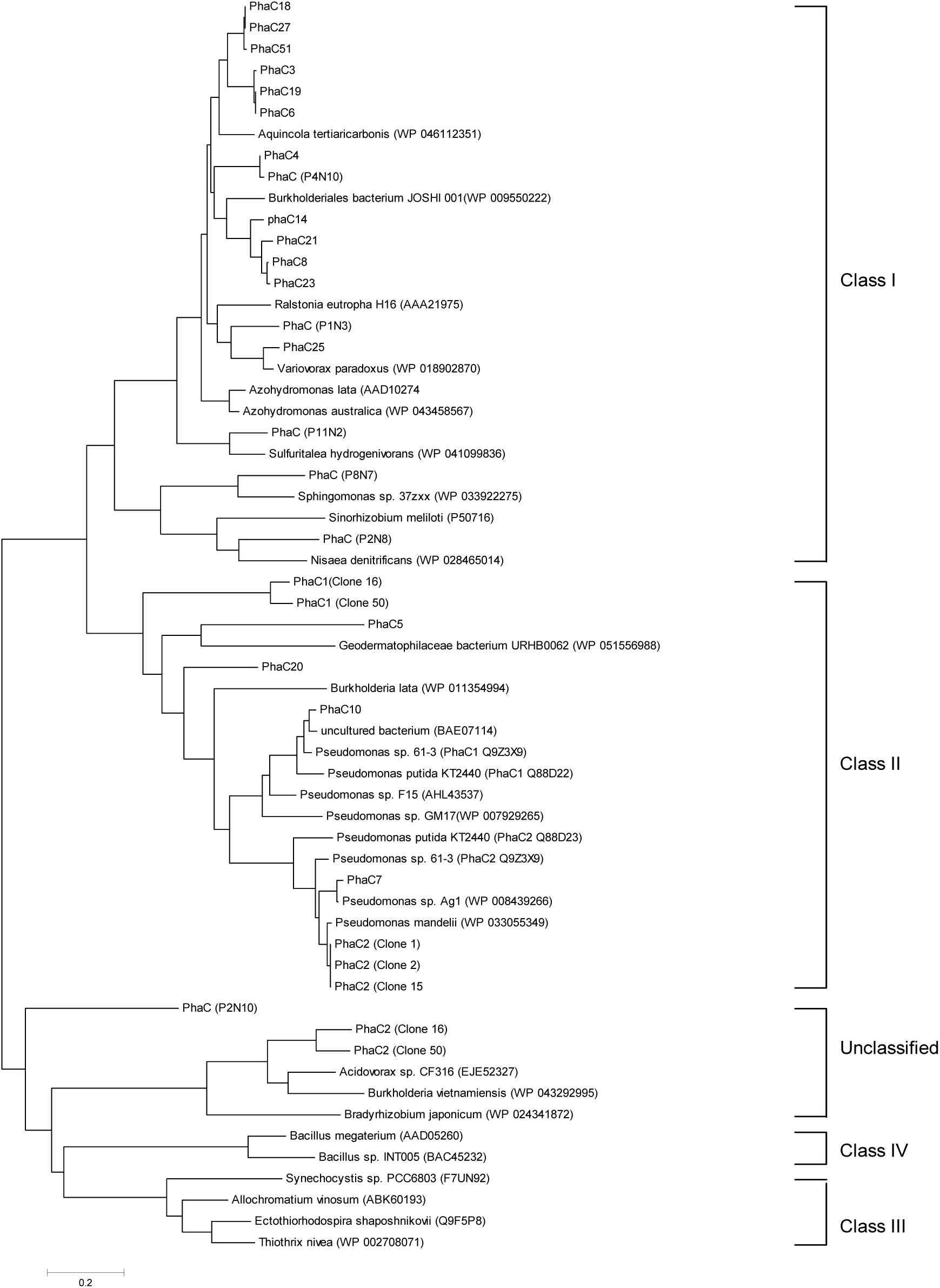
Phylogenetic analysis of the polyhydroalkanoate synthases (PhaC) of 11AW metagenomic library clones. Protein sequence alignments were performed using MUSCLE (Edgar 2004). Neighbour-jointing phylogenetic trees were generated with MEGA6 (Tamura et al. 2013). GenBank accession numbers of 11AW PhaC proteins: PhaC2_1_ (ALV86274), PhaC_3_ (ALV86289), PhaC_4_ (ALV86299), PhaC_5_ (ALV86308), PhaC_6_ (ALV86351), PhaC_7_ (partial, ALV86358), PhaC_8_ (ALV86364), PhaC_10_ (partial, ALV86387), PhaC_14_ (ALV86397), PhaC1_16_ (ALV86417), PhaC2_16_ (ALV86419), PhaC_18_ (ALV86462), PhaC_19_ (ALV86476), PhaC_20_ (ALV86493), PhaC_21_ (ALV86517), PhaC_23_ (ALV86529), PhaC_25_ (ALV86574), PhaC_27_ (ALV86602), PhaC1_50_ (ALV86626), PhaC2_50_ (ALV86626), PhaC_51_ (ALV86651), PhaC_P1N3_ (ALV86715), PhaC_P2N8_ (ALV86750), PhaC_P2N10_ (partial, ALV86755), PhaC_P4N10_ (ALV86768), PhaC_P8N7_ (ALV86771).

Class II PHA genes are commonly clustered with *phaZ* flanked by two PHA synthase encoding genes, *phaC1* and *phaC2*. Nine clones carried Class II *phaC* genes (Table 3). The *pha* gene locus (*phaC1*_*1*_-*phaZ*-*phaC2*_*1*_-*phaD*-*phaF-phaI*) in clone 1 was similar to the canonical locus in *Pseudomonas* including *P. putida* KT2440 (de Eugenio et al. 2010), except the presence of *orf7* encoding a PHA granule associated protein (Pfam09650) (Fig. 3A). Clone 2 and clone 15 were identical to the clone 1, except that they contained an insertion of the 17,013-bp transposon Tn4652 at 13,740 nt and 16,362 nt respectively (Supplementary 2). The Tn4652 duplicated target sequences were AACTC in clone 2 and TAGGA in clone 15. The transposon insertions in these clones are in opposite orientation. The same Tn4652 is present in *P. putida* KT2440 genome (3,366,550 −3,383,562 nt, GenBank: AE015451), located distant from the *pha* gene operon. These insertions likely occurred following introduction of the cosmid clones into *P. putida*.

**Fig. 3.**
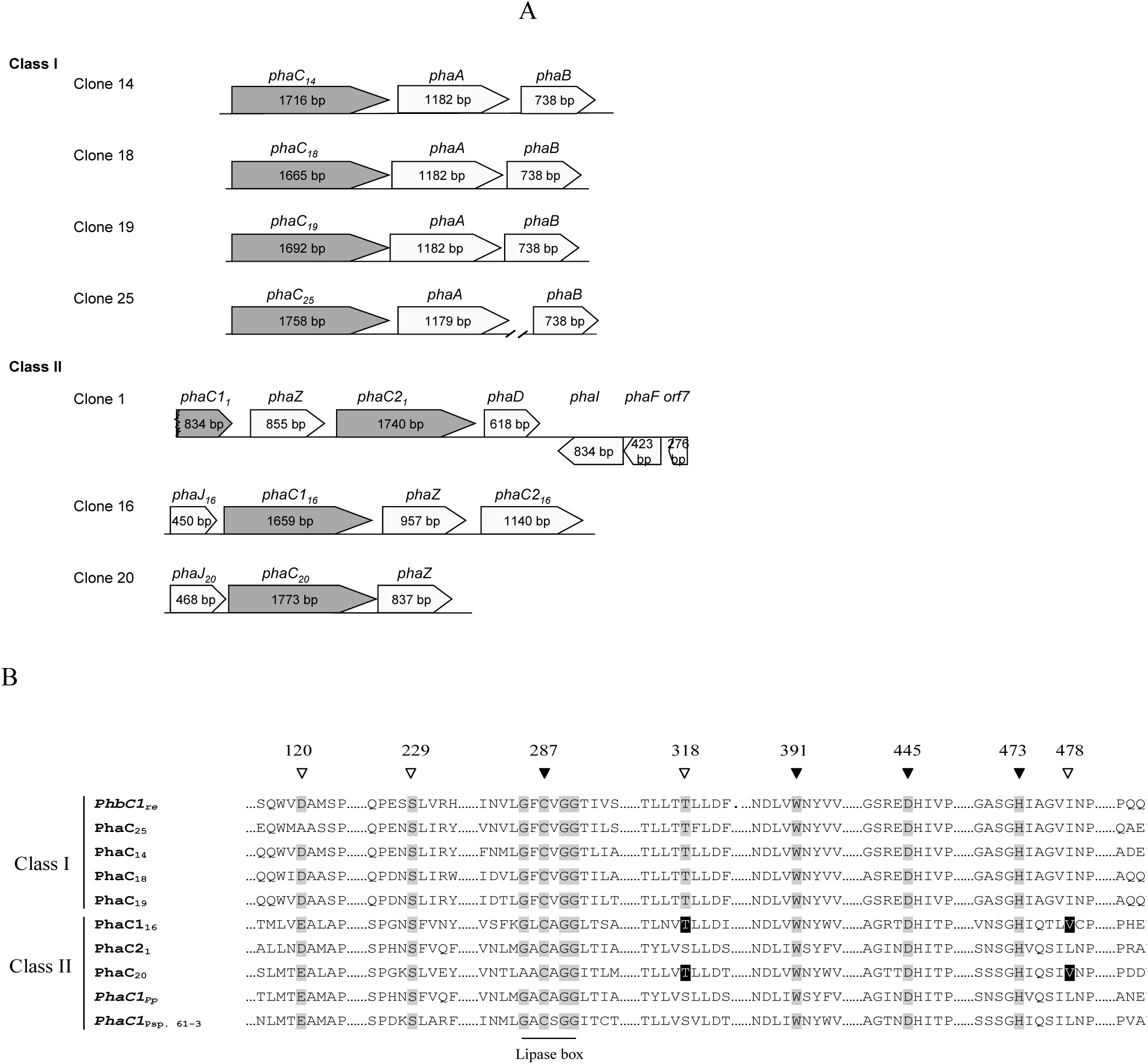
The *pha* genes and PhaC proteins in 11AW metagenomic DNA library clones. (A) The PHA^+^ clones are classified into Class I and II based on the PhaC protein sequences. (B) Conserved amino acids required for the activity of PHA synthases are indicated in closed triangles. Residues affecting substrate specificity were marked by open triangles. The positions of amino acid residues were numbered based on the sequence of PhaC1_16_ [GenBank: ALV86417]. GenBank accession numbers of PhaC proteins: PhaC_14_, ALV86397; PhaC_18_, ALV86462; PhaC_19_, ALV86476; PhaC_25_, ALV86574; PhaC1_2_, ALV86274; PhaC_20_, ALV86493; PhaC1_16_, ALV86417; *C. necator* PhbC1, AAA21975; *P. putida* PhaC1, Q88D25; and *P*. sp. 61-3 PhaC1, BAA36200.

Class II clones 5, 16, 20 and 50 had no genes encoding *phaD* or *phaFI* homologs. A *phaJ* gene encoding R-specific enoyl-CoA hydratase was identified in those clones (Fig. 3A; Supplementary 1). The *phaJ* gene was located immediately upstream of the *phaC* genes in clones 16, 20 and 50, but downstream of *phaC* in clone 5. In addition, a gene encoding PHA depolymerase (PhaZ) was located downstream of the *phaC* genes in clones 16, 20 and 50. ORFs downstream of the *phaZ* genes in clones 16 and 50 encoded proteins (PhaC2_16_ and PhaC2_50_) homologous to unclassified PHA synthases (Fig. 2; Supplementary 1). The proteins shared conserved regions in PhaC enzymes, but were ~30% shorter in N-terminal regions than in those of Class I/II PhaC (Supplementary 2).

Two ORFs encoding putative PhaC and PhaE proteins were identified in partially assembled clone P2N10 (Table 3; Supplementary 1). BLASTP showed the P2N10 PhaC protein best matched to the Class III PhaC [GenBank: WP_002708071] (43% identity) from γ-*Proteobacteria Thiothrix nivea* (Table 3). The available C-terminal sequence of P2N10 PhaE (234 amino acids) only exhibited 26% identity to the Class III PhaE [GenBank: WP_002708072] of *T. nivea*. In addition, the P2N10 PhaC protein was only 29% identical to the Class IV PhaC [GenBank: AAD05620] of *Bacillus megaterium*. Phylogenetic analysis suggested that P2N10 PHA synthase was distant from Class III/IV PhaC and might represent a new subclass of PHA synthases (Fig. 2).

Multiple alignment of amino acid sequences of each of the identified 11AW PhaC proteins showed the conserved catalytic triad (C287, D445 and H473 in PhaC1_16_), a tryptophan essential for dimerization (W391 in PhaC1_16_), and the lipase box GXCXGG (Jia et al. 2000) except that Ala replaced the first Gly residue in PhaC_20_ and second Gly in clone P8N7 (Fig. 3B; Supplementary 3). The serine residue (S229 in PhaC1_16_) required for PhaC activity (Hoppensack et al. 1999) was also conserved in all 11AW PhaC proteins. We chose the PhaC proteins in the Class I clones 14, 18 and 25, and the Class II clones 1, 16 and 20, for further study. We examined the nature of PHA produced using different carbon sources that result in substrate production either through fatty acid synthesis or β-oxidation.

### Class II clone 1 synthesizes MCL PHA

The cloned metagenomic DNA in clone 1 contained 42,747 bp (Table 3) [GenBank: KT944254] and most likely originated from *γ-Proteobacteria Pseudomonas*. The PhaC1_1_, PhaZ, PhaC2_1_ and PhaD proteins [GenBank: ALV86243, ALV86281, ALV86274 and ALV86275] were 82%, 92%, 73% and 78% identical to the corresponding orthologs (PhaC1, PhaZ, PhaC2 and PhaD) of *P. putida* KT2440. Clone 1 PhaC2_1_ was phylogenetically related to the PHA synthase 2 [GenBank: BAA36202] of *Pseudomonas* sp. 61-3 (Fig. 2). In contrast to other Class II PhaC, the conserved amino acid at position 129 of PhaC2_1_ was Asp rather than Glu (Fig. 3B). It has been previously demonstrated that substitution of the conserved Glu with Asp in Class II PhaC improves PHA yield with an increase of 3-hydroxybutyrate monomer (Matsumoto et al. 2005).

PHA synthesis in clone 1 was likely contributed solely by PhaC2_1_ because the 5’-region of *phaC1*_*1*_ was absent in the cloned DNA. Expression of the functional *phaZ* and *phaC2*_*1*_ might be driven by the promoters upstream of the individual genes, as occurs in *P. putida* KT2440 (de Eugenio et al. 2010). *P. putida* PpUW2 (PHA^-^) carrying clone 1 synthesized 3-hydroxyhexanote (3HHx, C6) and 3-hydroxyoctanoate (3HO, C8) copolymer of ~95% C8 monomer when grown with octanoate, similar to the PHA produced by wild-type *P. putida* PpUW1 (Table 4). When PpUW2 (clone 1) was grown with nonanoic acid, 3-hydroxynonanoate (3HN, C9) and 3-hydroxyheptanoate (3HP, C7) were incorporated into the PHA with greater 3HP than that synthesized by wild type PpUW1 (Table 4). These data suggest that those monomers were derived from β-oxidation of octanoate and nonanoic acid.

**Table 4.** Biosynthesis of SCL-and/or-MCL PHAs by 11AW metagenomic clones in *Pseudomonas putida*.

| *P. putida* | Carbon sources | PHA content (%) | PHA composition (mol%) |  |  |  |  |  |  |
| --- | --- | --- | --- | --- | --- | --- | --- | --- | --- |
|  |  |  | 3HB | 3HV | 3HHx | 3HP | 3HO | 3HN | 3HD |
| Class II |  |  |  |  |  |  |  |  |  |
| PpUW1 | Octanoate | 30.0 |  |  | 0.7 |  | 99.3 |  |  |
|  | Nonanoic acid | 45.0 |  |  |  | 19.1 |  | 80.9 |  |
|  | Gluconic acid | 1.7 |  |  |  |  | 21.1 |  | 76.8 |
| PpUW2 (clone 1) | Octanoate | 32.7 |  |  | 4.5 |  | 95.5 |  |  |
|  | Nonanoic acid | 23.4 |  |  |  | 42.0 |  | 58.1 |  |
|  | Gluconic acid | 8.2 |  |  |  |  | 20.4 |  | 79.6 |
| PpUW2 (clone 16) | Octanoate | 25.1 | 17.3 |  | 39.3 |  | 43.2 |  |  |
|  | Nonanoic acid | 46.5 | 3.4 | 40.9 |  | 55.4 |  |  |  |
|  | Gluconic acid | 4.5 |  |  | 47.6 |  | 52.4 |  |  |
| PpUW2 (pJC123) | Octanoate | 17.5 | 13.9 |  | 30.3 |  | 51.3 |  |  |
| PpUW2 (pJC157) | Octanoate | 32.5 | 14.5 |  | 33.9 |  | 49.3 |  |  |
| PpUW2 (clone 20) | Octanoate | 17.6 |  |  | 17.3 |  | 82.7 |  |  |
|  | Nonanoic acid | 45.0 |  | 7.5 |  | 50.2 |  | 42.3 |  |
|  | Gluconic acid | 3.9 |  |  | 9.9 |  | 37.4 |  | 52.7 |
| Class I |  |  |  |  |  |  |  |  |  |
| PpUW2 (clone 14) | Octanoate | 41.8 | 98.9 |  | 1.1 |  |  |  |  |
|  | Nonanoic acid | 59.4 | 6.8 | 93.2 |  |  |  |  |  |
|  | Gluconic acid | 81.2 | 97.9 |  | 0.8 |  | 1.3 |  |  |
| PpUW2 (clone 18) | Octanoate | 6.4 | 97.9 |  | 0.5 |  | 2.4 |  |  |
|  | Nonanoic acid | 17.3 | 32.8 | 67.2 |  |  |  |  |  |
|  | Gluconic acid | 30.7 | 100.0 |  |  |  |  |  |  |
| PpUW2 (clone 19) | Octanoate | 30.9 | 98.8 |  | 0.5 |  | 0.7 |  |  |
|  | Nonanoic acid | 38.1 | 20.9 | 79.2 |  |  |  |  |  |
|  | Gluconic acid | 12.3 | 100.0 |  |  |  |  |  |  |
| PpUW2 (PhaC25) | Octanoate | 15.6 | 95.0 |  | 2.9 |  | 2.1 |  |  |
|  | Nonanoic acid | 43.1 | 25.2 | 74.9 |  |  |  |  |  |
|  | Gluconic acid | 4.1 | 100.0 |  |  |  |  |  |  |

*P. putida* PpUW1 (PHA<sup>+</sup>) and PpUW2 (PHA<sup>-</sup>) carrying PHA<sup>+</sup> cosmids of 11AW metagenomic clones were grown at 30°C for 48 h in 0.1N M63 media supplemented with 0.5% octanoate, 0.5% nonanoic acid or 1% gluconic acid. PHA content was represented as relative to cell dry weight. Values were the averages of triplicate cultures. The standard deviations were <10% of the values. Abbreviations: 3HB, 3-hydroxybutyrate (C4); 3HHx, 3-hydroxyhexanoate (C6); 3HO, 3-hydroxyoctanoate (C8); 3HD, 3-hydroxydecanoate (C10).

When gluconic acid was used as the sole carbon source, 3HO and 3-hydroxyldecanoate (3HD, C10) copolymer was produced in PpUW2 (PHA^-^) carrying clone 1, very similar to the production of the parental strain PpUW1 (PHA^+^) (Table 4). The monomer composition of PHA was similar in both strains, but total PHA accumulation was ~4 fold higher in gluconic acid-grown PpUW2 with clone 1 (Table 4). These results suggest that Class II PhaC1_2_ was able to synthesize MCL (C6-C10) PHA.

### Class II clone 16 synthesizes SCL-MCL copolymer

Clone 16 contains 33,900-bp metagenomic DNA [GenBank: KT944263] probably originated from *β*-*Proteobacteria Methylibium* (Table 3). A *phaJ* gene (*phaJ*_*16*_) is predicted to encode a R-specific enoyl-CoA hydratase [GenBank: ALV86416] (Fig. 3A), which converts the β-oxidation intermediate 2-enoyl-CoA to (R)-3-hydroxyacyl-CoA used as PhaC substrate (Fukui et al. 1998). The PhaJ_16_ was phylogenetically related to the PhaJ4a_Re_ and PhaJ4b_Re_ of *Ralstonia eutropha* (Kawashima et al. 2012), and PhaJ4_Pp_ of *P. putida* (Sato et al. 2011) (Supplementary 4A). The amino acid residues Asp39 and His44 conserved at the active site of the dehydratases were identified in PhaJ_16_ (Supplementary 3B), except that the Ser residue was replaced by Pro71 in PhaJ_16_, same as in the PhaJ orthologs of *R. eutropha* and *P. putida* PhaJ4.

The three ORFs downstream of the *phaJ*_*16*_ gene encoded PhaC1_16_, PhaZ and PhaC2_16_ (Fig. 3A). PhaC1_16_ of Class II PHA synthase most closely matched the *Pseudomonas* sp. F15 PhaC1 [GenBank: WP_021487788], exhibiting 48% amino acid identity. PhaC1_16_ was also 42.5% and 43.4% identical to the PhaC1 and PhaC2 proteins of *P. putida* KT2440. Thr318 and Val478 in the PhaC1_16_ might favour the 3HB incorporation into PHA as occurred in the engineered PhaC proteins (Takase et al. 2003; Chen et al. 2014). Another putative PHA synthase encoded by the *phaC2*_*16*_ gene downstream of the *phaC1*_*16*_*phaZ* genes was homologous to unclassified PHA synthases (Fig. 2).

The quantity of PHA accumulated in *P. putida* UW2 carrying clone 16 was comparable to that of parental PpUW1 when grown with octanoate, nonanoic acid or gluconic acid (Table 4). In contrast to the MCL PHA (C6-C8) present in octanoate-grown PpUW2 (clone 1) or wild-type PpUW1, about 20% 3HB was detected in the PHA (C4-C6-C8) accumulated in PpUW2 carrying clone 16 under the same conditions (Table 4). When PpUW2 (clone 16) was grown with nonanoic acid (C9), 3-hydroxyheptanoate (3HP, C7), 3-hydroxyvalerate (3HV, C5) and 3HB (C4) were polymerized, which was in contrast to the 3HP-3HN PHA in nonanoic acid-grown PpUW with PhaC1 or wild type PpUW1. Additionally, equal amounts of C6 and C8 monomers were detected in the PHA from gluconic acid-grown PpUW2 with clone 16. However, PHA with C8 and C10 monomers was accumulated in PpUW2 (clone 1) and PpUW1 under the same growth conditions. These data implied that the PhaC1_16_ and/or PhaC2_16_ prefer substrates with shorter carbon chains than the PhaC2_1_ in clone 1 and both PhaC proteins of *P. putida* KT2440.

To further elucidate the activity of PhaC1_16_ and PhaC2_16_, the genes were cloned downstream of the constitutively active P*lac* promoter. When *P. putida* PpUW2 carrying pJC123 (P*lac*∷*phaC1*_*16*_) was grown with octanoate, SCL-MCL copolymer was of similar composition of C4-C6-C8 monomers to that synthesized by clone 16 (Table 4). However, PHA was not detected in strain PpUW2 carrying pJC159 (P*lac*∷*phaC2*_*16*_) under the same growth conditions. Coexpression of *phaC1*_*16*_ and *phaC2*_*16*_ in pJC157 (*Plac*∷*phaC1*_*16*_-*phaC2*_*16*_) in octanoate-grown PpUW2 resulted in synthesis of PHA with the similar monomer composition as those in the polymers produced in PpUW2 carrying clone 16 or pJC123 (P*lac*∷*phaC1*_*16*_) (Table 4). These data indicated that only the PhaC1_16_ was involved in SCL-MCL PHA biosynthesis in clone 16.

The *pha* gene locus of clone 16 was very similar to that of clone 50 (Supplementary 1), but the DNA in clone 50 probably originated from γ-*Proteobacteria Hydrocarboniphata* (Table 3). Both the Class II PhaC1 and unclassified PhaC2 in clones 16 and 50 shared 84% identity respectively. Expression of the *pha* genes was probably driven by a promoter upstream of *phaJ* genes.

### Class II clone 20 synthesizes MCL or SCL-MCL PHA

PHA^+^ clone 20 contained 30,964-bp metagenomic DNA [GenBank: KT944266], which probably originated from *Salinibacter* within the *Bacteroidetes* (Table 3). The cloned DNA contained only a single *phaC*_*20*_ gene (Fig. 3A). A putative fatty acid CoA ligase/ synthetase [GenBank: ALV86493] (ORF9; Supplementary 1) might be involved in degradation of fatty acids to acyl-CoA (the first step of β-oxidation cycle). The acyl dehydratase PhaJ_20_ [GenBank: ALV86492] was 56% identical to the PhaJ_16_. Conserved amino acid residues Asp41 and His46 at the active sites of the acyl dehydratases were present (Supplementary 4B) except that the Ser residue was replaced by Pro73 in the PhaJ_20_ protein. The *phaJ*_*20*_, *phaC*_*20*_ and *phaZ* genes in clone 20 were similar to those of the corresponding genes in clones 16 and 50 (Fig. 3A; Supplementary 1). In addition, the *pha* gene cluster of clone 20 was similar to that in *Burkholderia* sp. 383 (1,576,365 −1,581,635 nt) [GenBank: NC_007511], but a gene encoding a phasin protein is located between the *phaC* and *phaZ* genes in the *Burkholderia* genome.

The single PhaC_20_ [GenBank: ALV86493] best matched to its ortholog from *Burkholderia lata* at 59% amino acid sequence identity [GenBank: WP_011354994]. PhaC_20_ was 46-53% identical to the PhaC1_16_, PhaC2 and PhaC_5_ as well as PhaC1 and PhaC2 of *P. putida* KT2440. Presence of amino acid residues Thr353 and Val516 in PhaC_20_ (Fig. 3B) might favour substrates with fewer carbons than those preferred by clone 1 and *P. putida* PhaC proteins, as previously demonstrated with engineered Class II PHA synthases (Takase et al. 2003; Chen et al. 2014).

Clone 20 was able to synthesize 3HHx-3HO copolymer (C6-C8) with octanoate (Table 4), as occurred in PpUW2 (clone 1) and *P. putida* wild type. However, the proportion of C6 monomer increased 4-35 fold in the PHA isolated from PpUW2 (clone 20), compared to that in PpUW2 (clone 1) and wild-type PpUW1 (Table 4). When nonanoic acid was supplied, PpUW2 (clone 20) produced PHA with 8% 3-hydroxyvalerate (3HV), which was absent in the PHA synthesized by clone 1 and PpUW1 (Table 4). In contrast to the SCL-MCL PHA produced by clone 16, 3HB was not detected in the nonanoic acid or octanoate-grown PpUW2 with clone 20.

In contrast to the absence of detectable C6 monomer in the PHA produced in PpUW2 with clone 1 or *P. putida* wild type, 3HHx (~10%) was incorporated into the MCL PHA (C6-C8-C10) in PpUW2 (clone 20) (Table 4) when gluconic acid was used as the sole carbon source. About half the quantity of monomers was C10 in the PHA accumulated in PpUW2 (clone 20) grown with gluconic acid. These results suggested that the carbon chain lengths of favourite monomers of PhaC_20_ were between those of PhaC1_16_ and PhaC2_1_.

### Class I clone 14 synthesizes SCL-MCL PHA

The DNA in clone 14 probably originated from *β-Proteobacteria Methylibium* (Table 3). The PhaC_14_, PhaA and PhaB [GenBank: ALV86397, ALV86398 and ALV86399] were 65%, 76% and 74% identical to the PhbC1 (A16_1437), PhbA (A16_1438) and PhbB1 (A16_1439) of *C. necator* H16, respectively. In addition, PhaC_14_ was 48% identical to the PhbC protein (Smc002960) of *S. meliloti* Rm1021. The best match of PhaC_14_ was the PhaC protein of *Burkholderia* JOSHI_001 at 74% (Table 3). The *phaC*_*14*_*AB* locus of clone 14 is similar to that of clones 8, 21 and 23 (Supplementary 1). Expression of the *phaC*_*14*_, *phaA* and *phaB* genes in clone 14 was most likely driven by a promoter upstream of the *orf9* encoding a hypothetical protein.

When PpUW2 (clone 14) was grown with nonanoic acid, 7% 3HB was incorporated into 3HB-3HV copolymer (Table 4). 3HB was the primary monomer of PHA in octanoate-or gluconic acid-grown PpUW2. Both C6 and C8 monomer were present in the PHA with gluconic acid as carbon source. However, C8 monomer was absent in the PHA when octanoate was supplied (Table 4). *P. putida* PpUW2 carrying clone 14 produced the highest quantity of PHA among all the PHA^+^ clones (Table 4). These results indicated that PhaC_14_ could synthesize SCL-MCL copolymer though 3HB was the dominant monomer.

### Class I clone 18 synthesizes SCL and SCL-MCL PHA

Clone 18 contained 37,818-bp metagenomic DNA [GenBank: KT944264], which probably originated from *β-Proteobacteria Rubrivirax* (Table 3). The annotated *phaC*_*18*_, *phaA* and *phaB* were organized in one operon (Fig. 3A), as also occurred in clones 27 and 51 (Supplementary 1). PhaC_18_ [GenBank: ALV86462] shared 100% and 99% identity to PhaC_27_ [GenBank: ALV86602] and PhaC_51_ [GenBank: ALV86651] respectively. The PhaC_18_ protein best matched to its homolog in *Azohydromonas austrilica* (Table 3), and was 71%, 49% and 61% identical to the PhaC_14_, PhbC_Sm_ and PhbC1_Re_ respectively.

SCL-MCL PHA (C4-C6-C8) was produced when PpUW2 (clone 18) was grown with octanoate (Table 4), as occurred in PpUW2 with clone 16. However, the fraction of C6-C8 monomers was only 3% in the PHA from clone 18, in contrast to the ~83% in the copolymer from clone 16. The yield of PHA was ~7 fold less than that produced by clone 14, but C8 monomer was absent (Table 4). PHA of 3HB-3HV monomers was accumulated in nonanoic acid-grown PpUW2 (clone 18). The ratio of 3HB to 3HV increased 5 fold compared to that in PpUW2 (clone 14) under the same growth conditions (Table 4). When PpUW2 carrying clone 18 was grown on gluconic acid, only PHB was produced. The yield of polymer increased 5 fold compared to that in octanoate-grown cells, and also the second highest quantity of PHA produced among all strains grown on gluconic acid. These results implied that PhaC_18_ could synthesize SCL-and SCL-MCL PHA dependent of the available carbon source.

### Class I clone 19 synthesizes SCL and SCL-MCL PHA

Clone 19 contained 25,085-bp of DNA that might originate from *β-Protoebacteria Leptothrix* (Table 3) [GenBank: KT944265]. Three ORFs encoded PhaC_19_, PhaA and PhaB for PHA biosynthesis (Fig. 3A). PhaC_19_ shared 99% identity with PhaC_6_ and PhaC_3_ (Fig. 2). The best match for PhaC_19_ is the ortholog of *A. austrilica* at 73% (Table 3). The *phaC*_*19*_*AB* genes in clone 19 probably consisted of one operon, and the promoter was located upstream of the *phaC*_*19*_.

Clone 19 was able to synthesize SCL-MCL PHA (C4-C6-C8) with octanoate but only PHB (C4) in gluconic acid-grown PpUW2 (Table 4), as occurred in PpUW2 carrying clone 18. However ~5 fold more PHA was synthesized by clone 19 than was produced by clone 18 in octanoate medium (Table 4). In contrast, the quantity of PHA was lower in gluconic acid-grown PpUW2 with clone 19 than it was in the same host carrying clone 18. Moreover, PHA composed of 3HB-3HV monomers was synthesized in noanoic acid-grown PpUW2 (clone 19), as occurred in PpUW2 (clone 18). These data suggested that PhaC_19_ was able to synthesize both SCL and SCL-MCL PHA.

### Class I clone 25 synthesizes SCL and SCL-MCL PHA

Clone 25 contained 41,012-bp of DNA [GenBank: KT944269], which was predicted to originate from *β-Proteobacteria Variovorax* (Table 3). The *phaC*_*25*_ and *phaA* genes were apparently located in a single operon (Fig. 3A). A *phaB* gene was located 9 open reading frames downstream of the *phaA*. The *phaC*_*25*_*A* and *phaB* loci in clone 25 are similar to those in clone P1N3 (Supplementary 1), though the *phaB* gene in clone P1N3 was four open reading frames away from *phaC*_*P1N3*_*A*. PhaC proteins of clones P1N3 and 25 shared 77% identity. In addition, PhaC_25_ was 35% and 61% identical to PhbC_Sm_ and PhbC1_Re_. The conserved amino acid Asp130 affecting PhaC substrate specificity (Matsumoto et al. 2005) was replaced by Ala152 (Fig. 3B).

SCL-MCL PHA (C4-C6-C8) was produced in PpUW2 harbouring clone 25 when grown on octanoate, and only PHB (C4) was isolated from gluconic acid-grown cells (Table 4). Similar results were obtained in PpUW2 with clone 18 or clone 19 under the same conditions. However, the quantity of C6-C8 monomers was greater with clone 25 that those with clone 18 or 19. Nonanoic acid-grown PpUW2 (clone 25) produced 3HB-3HV copolymer similar to that from PpUW2 with clone 18 or clone 19 (Table 4). These data suggested that PhaC_25_ had similar properties to those of PhaC_18_ and PhaC_29_, but different from that of PhaC_14_.

### Complementation of *S. meliloti phaC* mutant with 11AW PHA^+^ clones

In an earlier study, *S. meliloti* Rm11476 (*exoY*∷Tn5 *phaC*∷Tn5-233) had been employed for functionally harvesting metagenomic library clones pCX92, pCX9M1, pCX9M3 and pCX9M5 expressing Class I PHA synthases (Schallmey et al. 2011). The PhaA/PhbB and PhaB/PhbB proteins encoded by the cloned metagenomic DNA and/or *S. meliloti* genome could supply (R)-3-hydroxybutyrate for PHB biosynthesis catalyzed by the PhaC/PhbC enzymes. In order to examine the possibility that the PHA^+^ clones of 11AW metagenomic library were able to complement the PHA^-^ phenotype of *S. meliloti* Rm11476, The Class I clones 14, 18, 19 and 25 (Table 3) were introduced into *S. meliloti* Rm11476. In each case, complementation was observed, indicating PhaC function in the *S. meliloti* background (Fig. 4), with greatest accumulation of PHA observed with clone 14. However, when clones pCX92, pCX9M1, pCX9M3 and pCX9M5 were transferred to *P. putida* PpUW2 (PHA^-^), PHA was not detected in the recombinant strains grown with either octanoate or gluconic acid (data not shown). These results strengthen the argument that multiple hosts should be employed to screen for novel *phaC* genes.

**Fig. 4.**
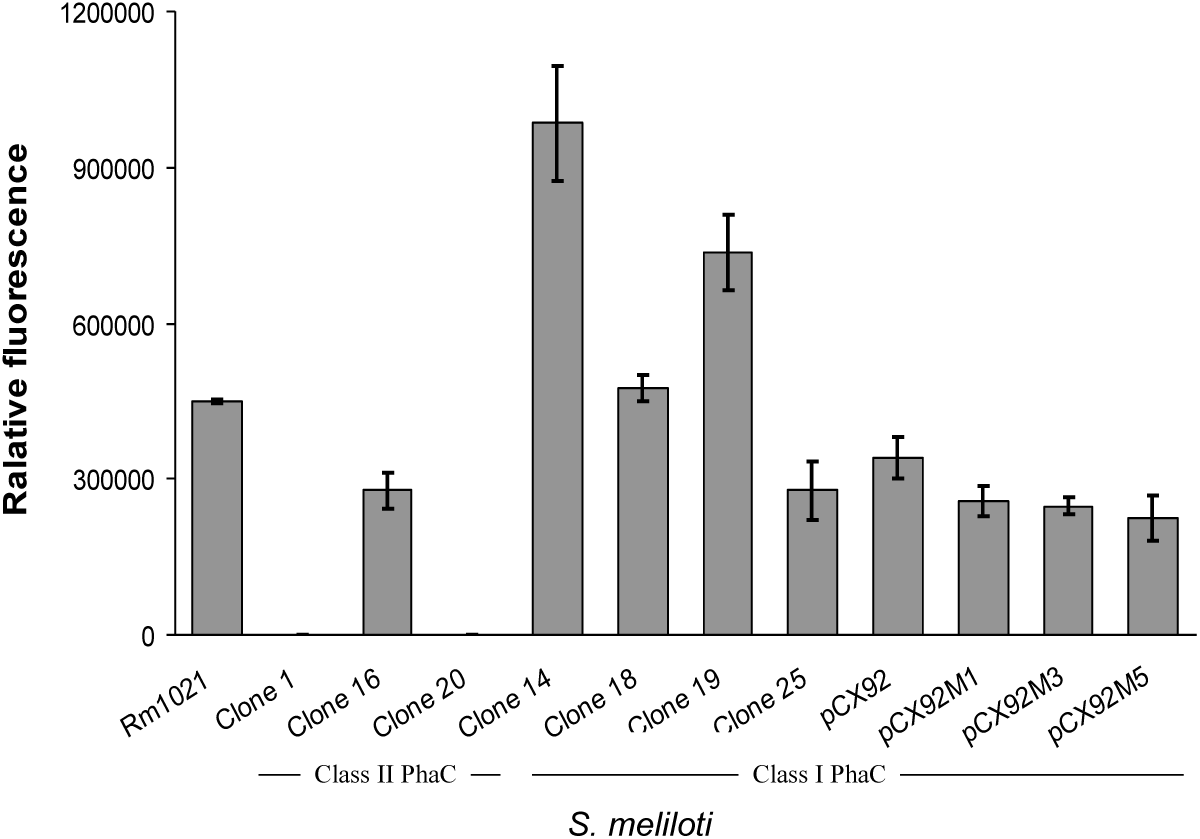
Complementation of *S. meliloti* Rm11476 (*phbC*) with metagenomic clones. The cosmid DNA of PHA^+^ clones encoding Class I and II PHA synthases was transferred to *S. meliloti* Rm11476 by conjugation. Recombinant strains were grown in YM medium. PHA production was estimated by measuring the fluorescence of Nile red stained cells. *S. meliloti* wild type Rm1021 and previously isolated Class I clones (CX92, pCX92M1, pCX92M3, and pCX92M5; (Schallmey et al. 2011)) were used as controls.

We then asked whether Class II *phaC* could function in the *S. meliloti* background. First, we observed no detection of PHA when *phaC1*_*Pp*_ (P*lac*∷*phaC1*_*Pp*_) was expressed in Rm11476 even though the *lac* promoter is functional and 3-hydroxybutyrate is supplied by the PhbAB_Sm_. Similarly, PHA^+^ clones 1 and 20 carrying Class II *phaC* genes also failed to complement the PHB^-^ phenotype of *S. meliloti* Rm11476 (Fig. 4), implying that the *phaC2*_*1*_ and *phaC*_*20*_ genes were not expressed and/or the substrate (R)-3-hydroxyalkanoic acids were absent in *S. meliloti*. However, when clone 16 was introduced into *S. meliloti* Rm111476, PHA was accumulated to a similar level as with Rm11476 containing PHB^+^ clones pCX92, pCX9M1, pCX9M3 or pCX9M5 (Fig. 4). These data indicated that the *phaC1*_*16*_ gene was expressed and its product was functional. These results suggest that *P. putida* is a more permissive surrogate host than *S. meliloti* for screening of novel PHA synthases.

## Discussion

Biosynthesis of SCL-MCL PHA requires PHA synthases having a broad range of substrate specificity and SCL and MCL precursors (R-3-hydroxyacyl-CoAs) must be available within the cell. Building on earlier work that has sought to mine metagenomic DNA for modification of bacterial PHA production, we have successfully isolated cosmid clones that are able to functionally complement a *P. putida* PHA synthesis mutant. DNA sequence analysis revealed that the isolated metagenomic DNA originated from a broad diversity of bacteria, and encoded either Class I or Class II PHA synthases enzymes. Of note is that DNA from most of the isolated clones did not very closely match known sequences. Also of interest is the influence that the clones had on the quality and quantity of PHA produced in the *P. putida* surrogate host background.

A major appeal of PHA is that, as alternatives to conventional fossil fuel-derived plastics, they can be produced from renewable resources. The potential to improve the efficiency and cost of production, and expand the range of polymers and copolymers that are available for production, should impact on competitiveness and adoption of these materials in the marketplace. In this context, it is important to understand how the properties of PHA are determined by the PHA synthase enzyme and the substrate that is available to those enzymes, which is in turn influenced by the culture conditions and the metabolic pathways leading to substrate formation (Meng et al. 2014). PHA consisting of both short-chain-length (SCL, ≤C5) and medium-chain-length (MCL, ≥C6) monomers have great flexibility, decreased breakage and reduced melting point (Noda et al. 2005). Most naturally occurring PhaC proteins polymerize either SCL-or MCL-monomers, but there are a few know examples such as the Class I PhaC of *Aeromonas caviae* (Fukui and Doi 1997) and some *Pseudomonas* Class II PhaC (Matsusaki et al. 1998) that can synthesize SCL-MCL copolymers.

In the present work, heterologous complementation of the *P. putida* PHA^-^ strain facilitated efficient simultaneous screening of millions of 11AW metagenomic clones, resulting in recovery of a greater number of novel Class I, Class II, and unclassified PHA synthases than from previous reports (Schallmey et al. 2011; Cheema et al. 2012). Additionally, the clones carrying Class I *phaC* isolated in *P. putida* were able to complement the PHA^-^ phenotype of *S. meliloti*, but the previously isolated Class I clones isolated in *S. meliloti* (Schallmey et al. 2011) failed to synthesize PHA in *P. putida*. These results support the development of multi-host systems to increase the chances of the successful expression of *pha* genes of interest.

Biosynthesis of SCL-MCL PHA of various forms by Class I clones 14, 18, 19 and 25 and Class II clones 16 and 20 suggested availability of a full range of R-3-hydroxyacyl-CoAs derived from β-oxidation of fatty acids such as nonanoic acid and octanoate. The differential monomer compositions in PHA synthesized by the Class II enzymes PhaC_16_ and PhaC_20_ are of particular interest, especially since most Class II PHA synthases are unable to incorporate SCL monomers such as 3HB and 3HV. Changes of conserved amino acid residues Ser325Thr, Leu484Val, and Glu508Leu (conserved positions in *P. putida* PhaC1) have been identified to be involved in PhaC substrate specificity and PHA yield (Takase et al. 2003; Shozui et al. 2010; Chen et al. 2014). Presence of the amino acids Thr353 (corresponding to *P. putida* PhaC1 position 325), Val516 (corresponding to *P. putida* PhaC1 position 484) and Ala540 (corresponding to *P. putida* PhaC1 position 508) PhaC_20_ might contribute to the enzyme’s ability to incorporate SCL monomers compared to those of 11AW PhaC2_1_ and PHA synthases of *P putida* KT2440. Similarly, the presence of three residues (Thr317, Val478 and Leu502) at these positions in PhaC1_16_, most likely resulted in increased 3HB content as demonstrated previously with engineered Class II PHA synthases (Takase et al. 2003; Shozui et al. 2010; Chen et al. 2014).

## Conclusions

We obtained 27 clones encoding Class I, II and unclassified PHA synthases by functional screening of soil metagenomic cosmid clones in a *P. putida* PHA^-^ strain. Seven clones that were characterized in more detail were able to produce a broad range of polymers and copolymers, including SCL-MCL mixtures, depending on carbon source. Through this work we have demonstrated the potential for using metagenome-derived clones for production of a variety of PHAs of possible industrial utility. The collection of PHA metabolism genes from uncultivated organisms provides not only a resource for production strain development, but also a series of sequence templates that could prove useful in enzyme engineering efforts directed towards generation of PHA products with desired properties.

## Nucleotide sequence accession numbers

Complete sequences of 11AW metagenomic PHA^+^ DNA have been deposited in GenBank nucleotide sequence database [GenBank: KT944254-KT944278, KU728995 and KU728996].

## Abbreviations

*P. putida*: *Pseudomonas putida*
PHA: polyhydroxyalkanoate
PhaA/PhbA: β-ketothiolase
PhaB/PhbB: acetoacetyl-CoA reductase
PhaC/PhbC: PHA synthase
PhaZ: PHA depolymerase
3HA: 3-hydroxyalkanoate
3HB: 3-hydroxybutyrate
3HV: 3-hydroxyvalerate
3HHx: 3-hydroxyhexanoate
3HP: 3-hydroxyheptanoate
3HO: 3-hydroxyoctanoate
3HN: 3-hydroxynonanoate
3HD: 3-hydroxydecanoate
SCL: short chain length
MCL: medium chain length
*S. meliloti*: *Sinorhizobium meliloti*
GC-MS: gas chromatography-mass spectrometry
CDW: cell dry weight

## Authors’s contributions

JC and TCC conceived and designed this study, carried out all the experiments, and prepared the manuscript. Both authors read and approved the final manuscript.

## Acknowledgements

We are very grateful to Ricardo Nordeste for assisting with GC-MS analysis of PHA. We wish to thank Dr. Bruce Ramsay (Polyferm Canada) for providing PHA standards, and Dr. Josh Neufeld for use of the FilterMax F5 Multi-mode microplate reader.

## Ethical statement

The authors certify that this manuscript has not been published previously, and not under consideration for publication elsewhere, in whole or in part. No data have been fabricated or manipulated (including images), and no data, text, or theories by others are presented as if they were the authors’ own. Consent to submit has been received explicitly from all the authors listed. And authors whose names appear on the submission have contributed sufficiently to the scientific work and therefore share collective responsibility and accountability for the results. This article does not contain any studies with human participants or animals performed by any of the authors.

## Supplementary figures

**Supplementary 1.**
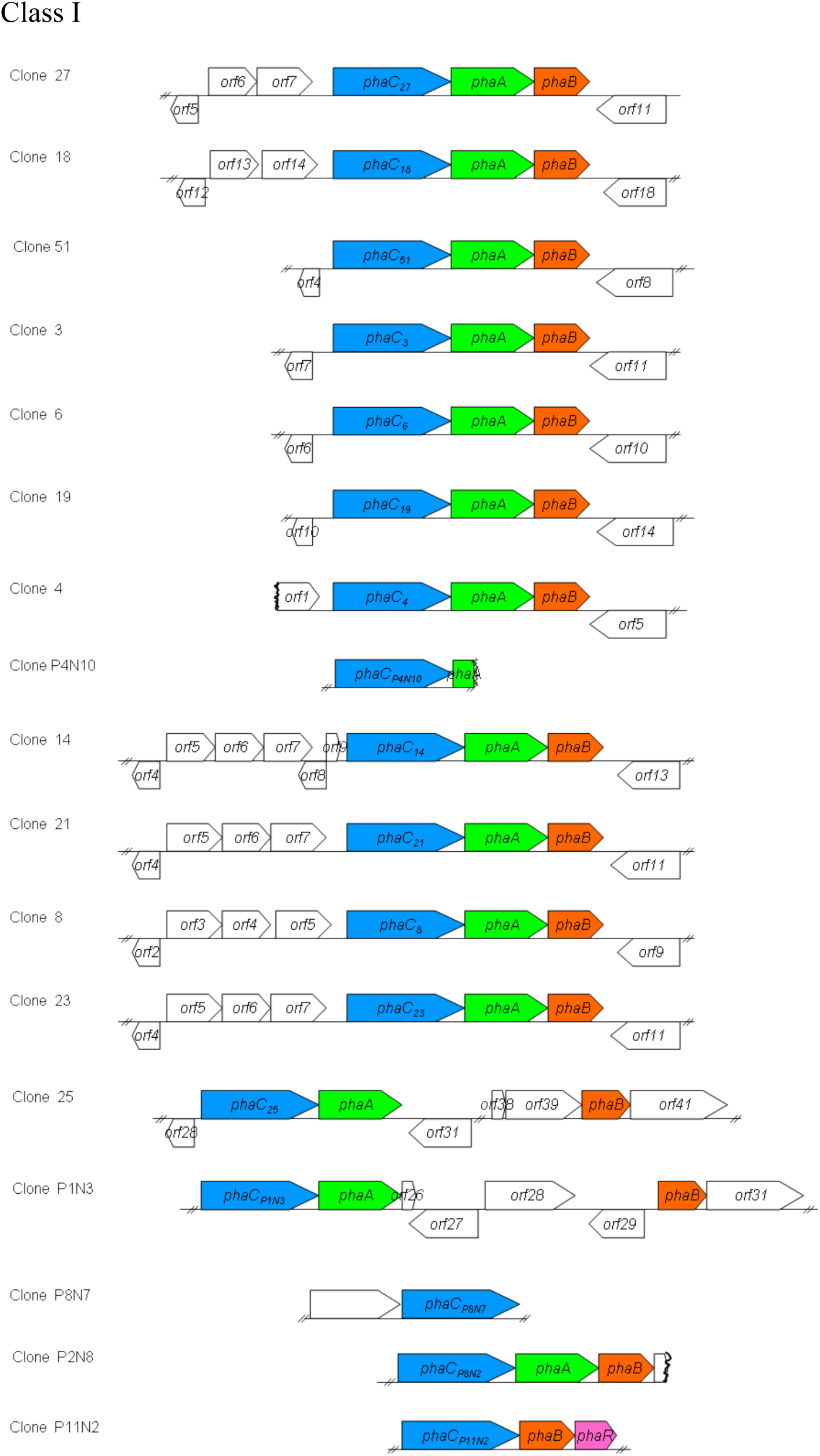
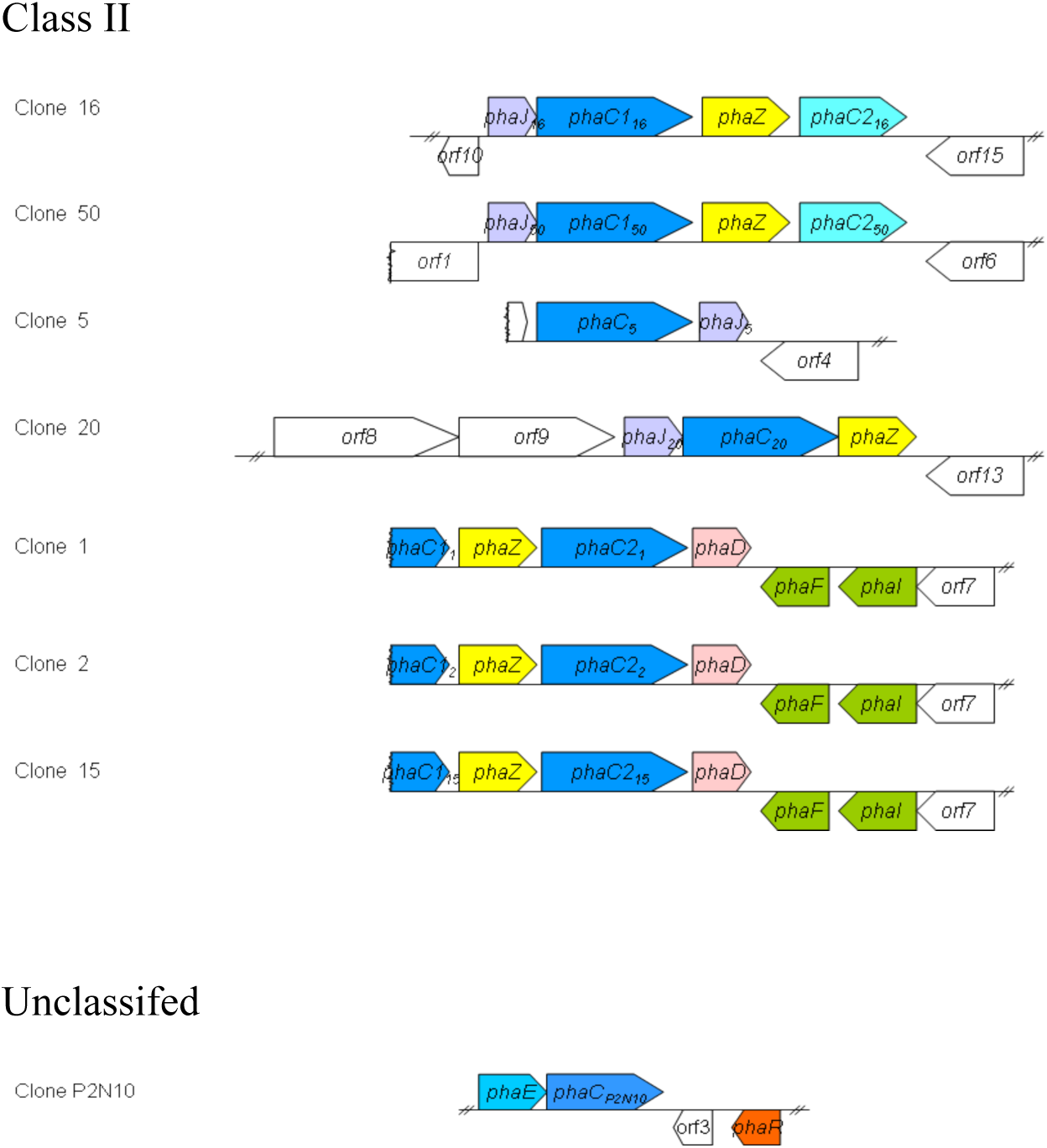
The *pha* gene locus in 11AW metagenomic DNA clones encoding PHA synthases (PhaC) were classified based on the PhaC protein sequences.

**Supplementary 2.**
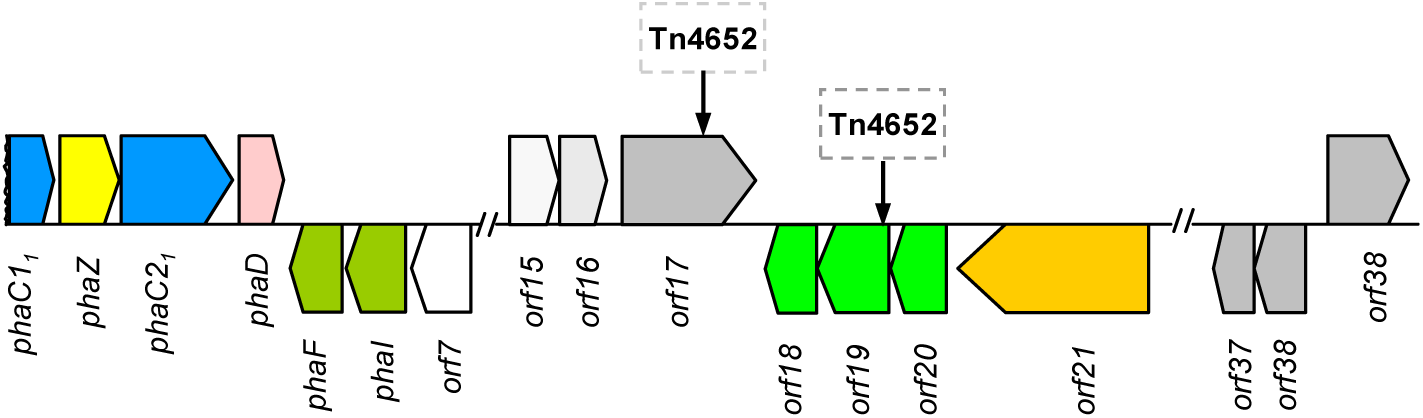
Transposon Tn4652 in 11AW PHA+ clones 2 and 15. Insertion of the Tn4652 into the *orf17* of clone 1 resulted in clone 2. Clone 15 was derived from clone 1 by the transposon insertion into the *orf19*.

**Supplementary 3.**
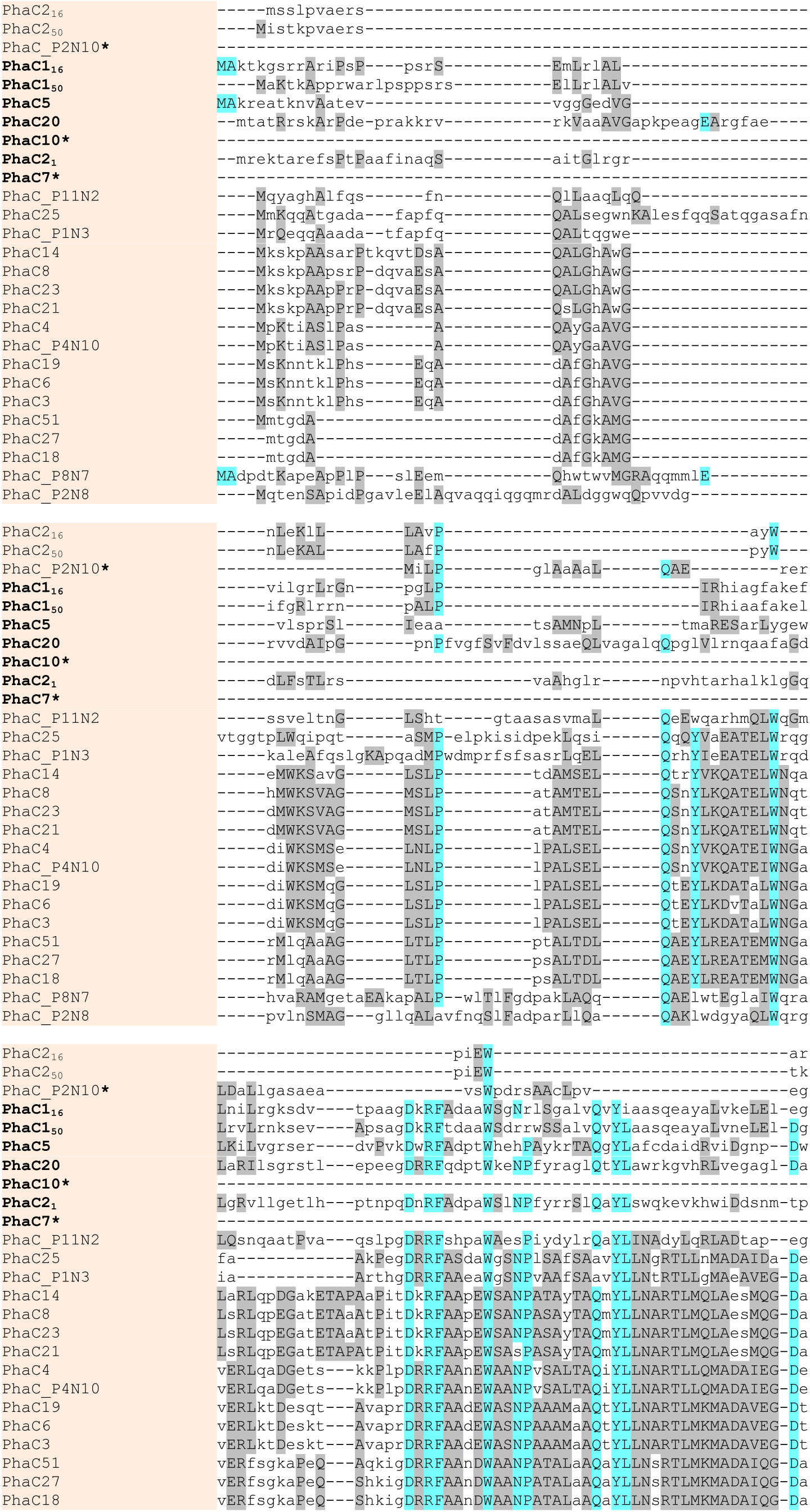
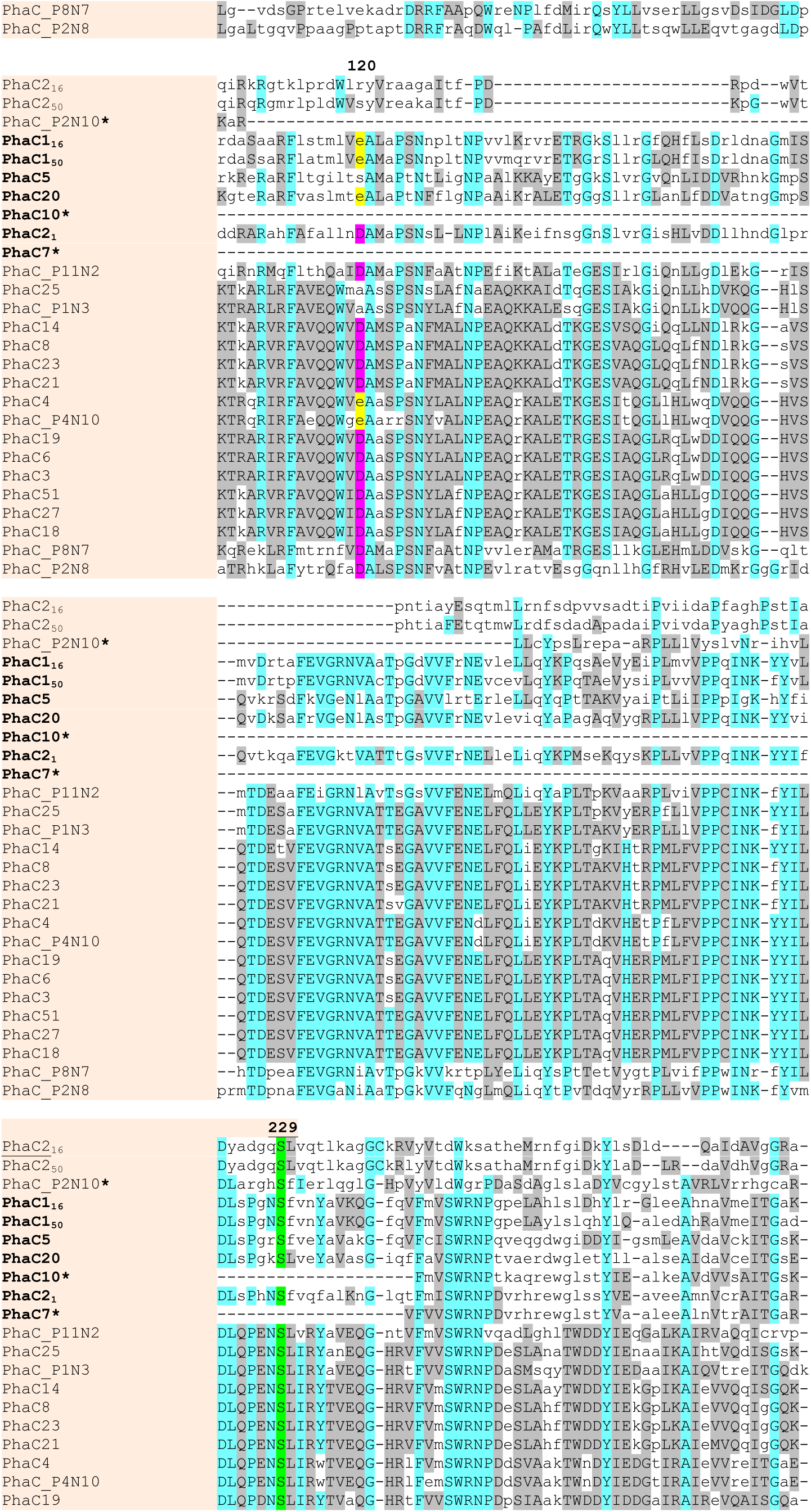
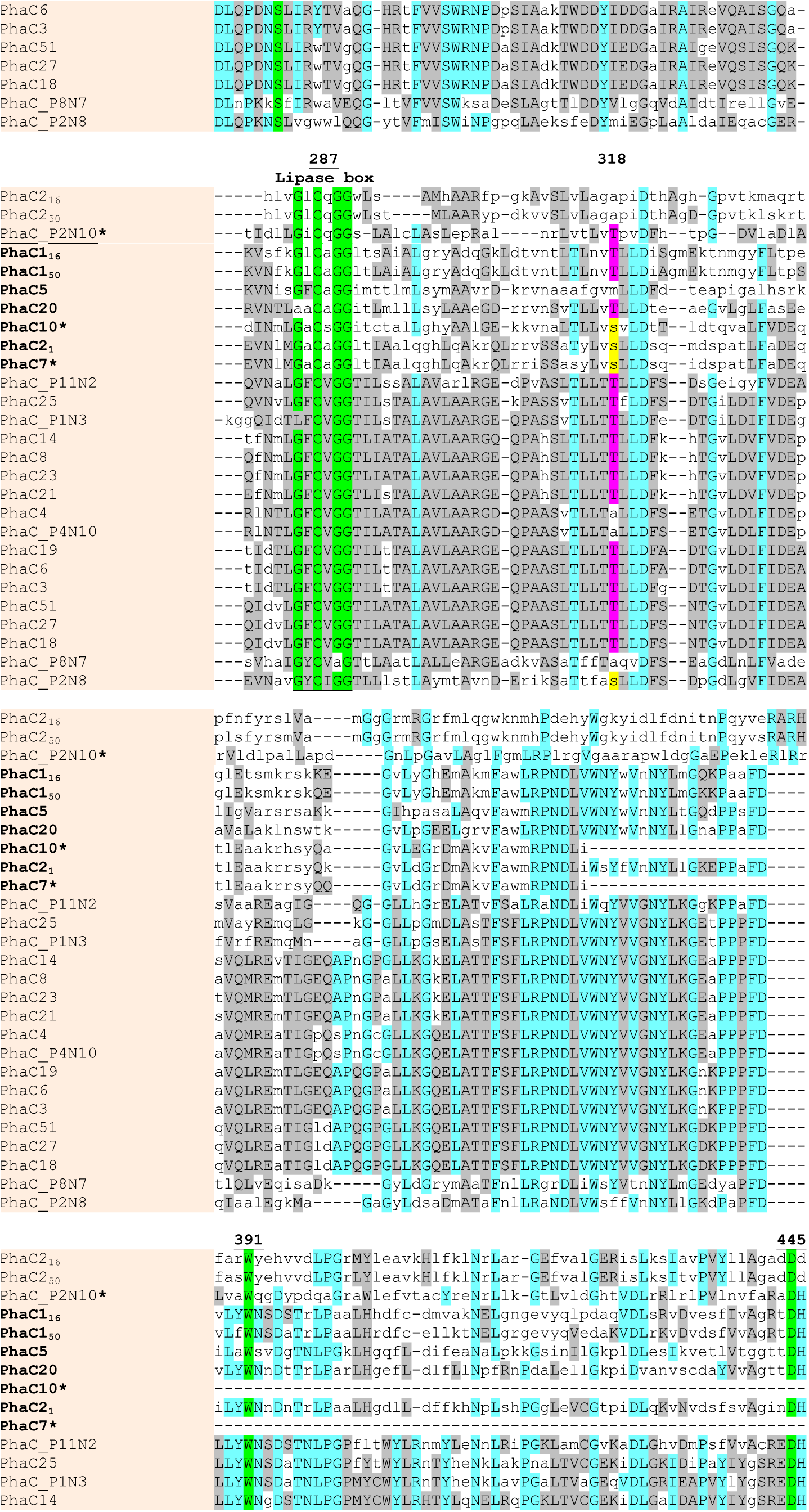
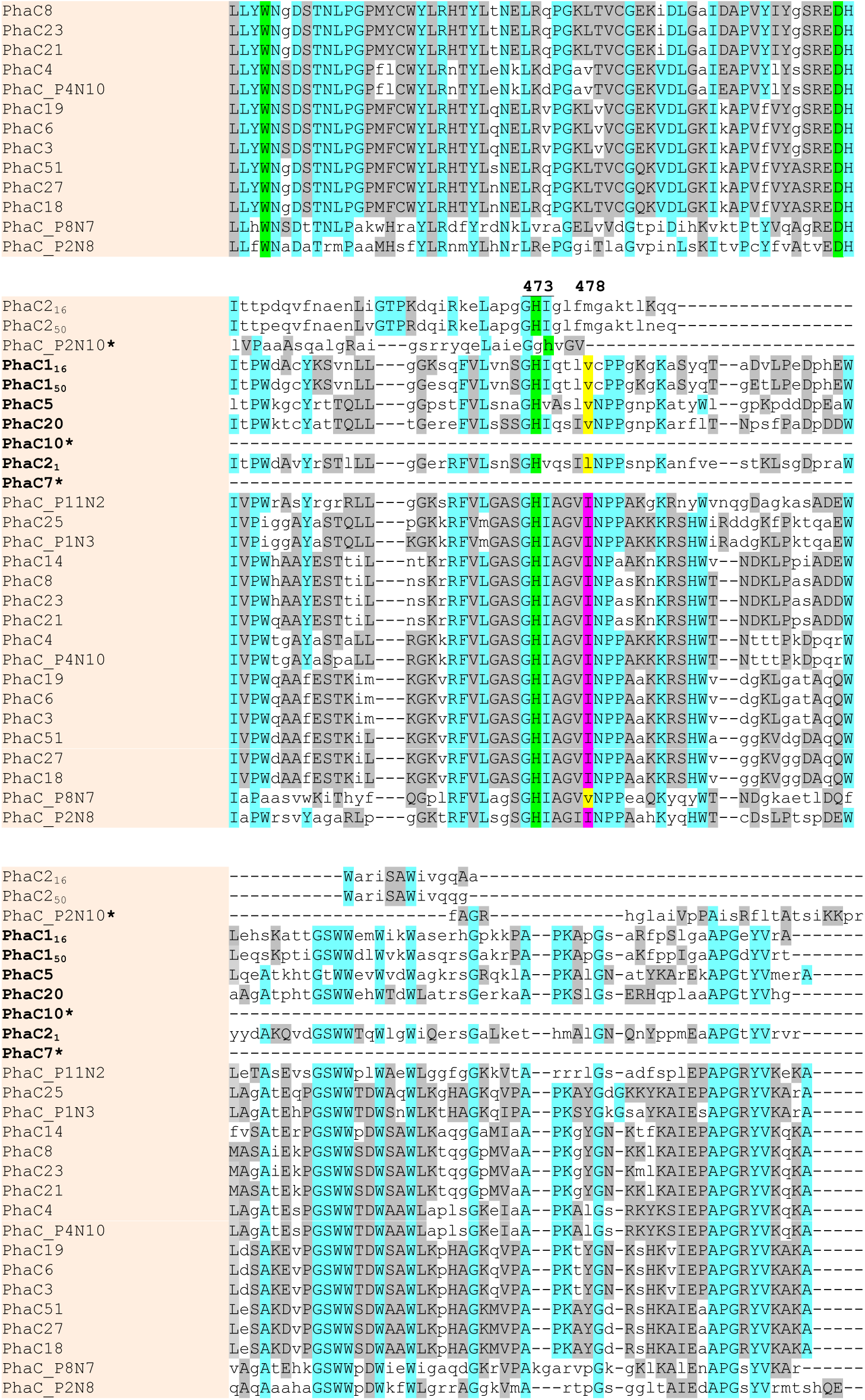
Multiple sequence alignment of PHA synthases of 11AW metagenomic DNA clones. Alignment was performed with MUSCLE. The conserved amino acid residues affecting substrate specificity in Class I and II PhaC are highlighted in purple and yellow respectively. The positions of amino acids essential for PhaC activity are highlighted in green. The position of amino acid residues are marked based on PhaC1_16_ sequence. Names of Class I PhaC are in regular font, Class II PhaC in bold, and unclassified PhaC underlined. PhaC proteins with partial sequences are marked with a star.

**Supplementary 4.**
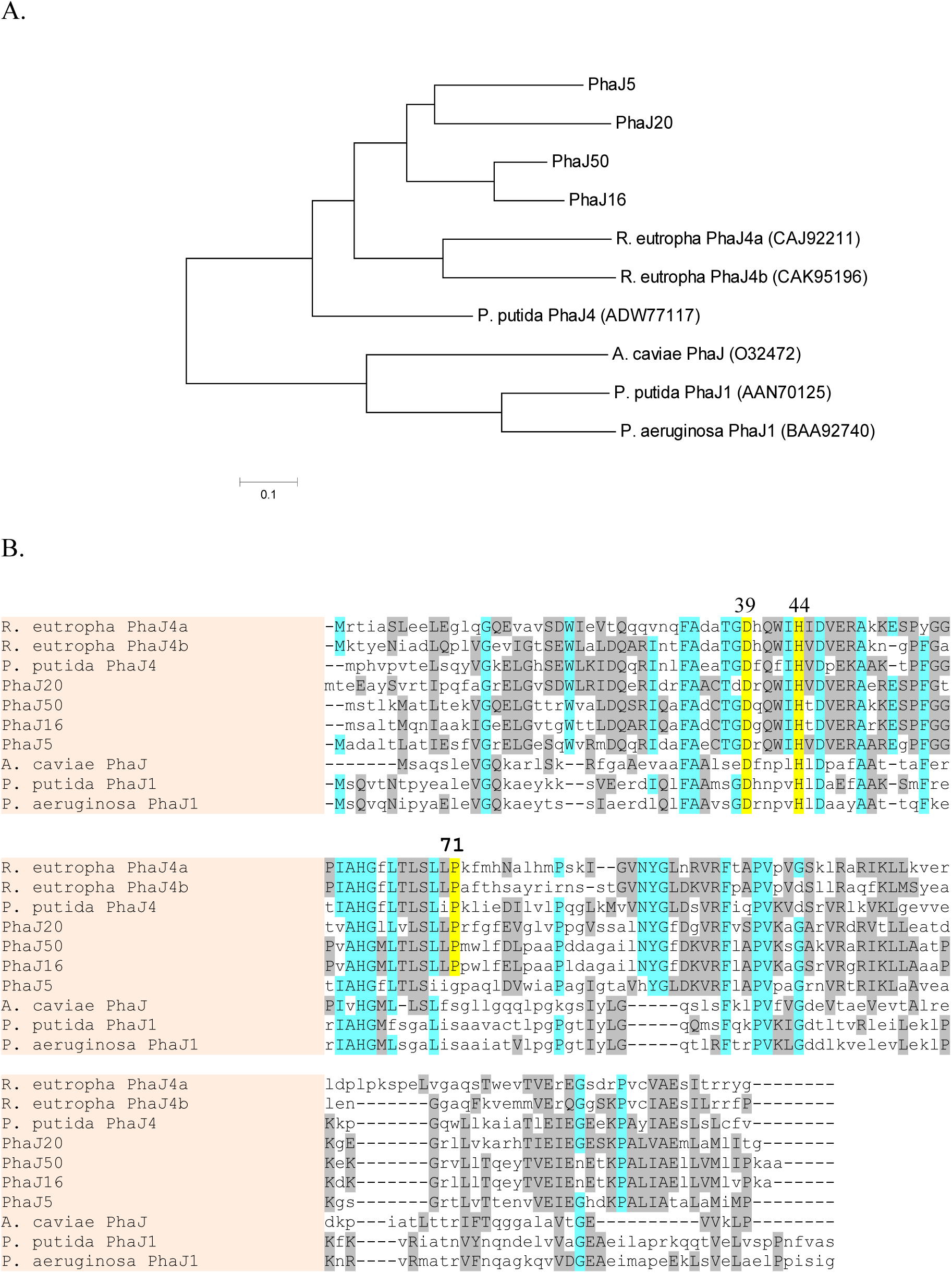
R-specific enoyl-CoA hydratases PhaJ. (A) Phylogenetic tree was constructed with MEGA6. The bar represents substitution of amino acid residue. (B) Conserved amino acid Asp^39^, His^44^ and Ser^7^ required for activity are marked.

